# 3D printing cytoskeletal networks: ROS-induced filament severing leads to surge in actin polymerization

**DOI:** 10.1101/2025.03.19.644260

**Authors:** Thomas Litschel, Dimitrios Vavylonis, David A. Weitz

**Affiliations:** School of Engineering and Applied Sciences, Harvard University, Cambridge, MA, USA; Department of Physics, Lehigh University, Bethlehem, PA, USA; Department of Physics, Harvard University, Cambridge, MA, USA; Wyss Institute for Biologically Inspired Engineering, Harvard University, Boston, MA, USA

## Abstract

The cytoskeletal protein actin forms a spatially organized biopolymer network that plays a central role in many cellular processes. Actin filaments continuously assemble and disassemble, enabling cells to rapidly reorganize their cytoskeleton. Filament severing accelerates actin turnover, as both polymerization and depolymerization rates depend on the number of free filament ends. Here, we use light to control actin severing in vitro by locally generating reactive oxygen species (ROS) with photosensitive molecules such as fluorophores. We see that ROS sever actin filaments, which increases actin polymerization in our experiments. However, beyond a certain threshold, excessive severing leads to the disassembly of actin networks, allowing us to selectively remove actin structures. Our experimental data is supported by simulations using a kinetic model of actin polymerization, which helps us understand the underlying dynamics. In cells, ROS are known to regulate the actin cytoskeleton, but the molecular mechanisms are poorly understood. Here we show that, in vitro, ROS directly affect actin reorganization.

---

Actin is the most abundant structural protein in eukaryotic cells, forming a dynamic filament network that undergoes continuous assembly and disassembly.^1^ This allows cells to rapidly reorganize their cytoskeleton in response to environmental and internal cues.

A key constraint on actin assembly is the formation of new filaments de novo. Actin polymerization is nucleation-limited, meaning that elongation of existing filaments is fast relative to the initiation of new filaments.^2^ Therefore, the amount of actin polymerization directly correlates with the number of existing filaments.

As nucleation from monomers is slow, filament severing offers an alternative mechanism to increase the amount of filaments. When a filament is severed, the number of filament ends doubles, therefore doubling polymerization sites. Severing can occur through various means; for example, mechanically^3-5^ or through interactions with certain enzymes.^6,7^

Here we focus on the potential role of reactive oxygen species (ROS), which are ubiquitous in cells and among the most reactive molecules in biological systems.^8,9^ ROS have been shown to greatly affect actin polymerization in vivo,^10-13^ however the underlying mechanisms are not well understood. In vitro, ROS are known to oxidize and damage actin monomers,^14^ which could cause filaments to break, thereby increase the number of polymerizing ends and affect the dynamics of actin assembly.

In this work we show that ROS interact with actin filaments and can increase actin polymerization drastically in vitro. We describe a simple experimental system that allows for the local generation of ROS via light-excitation of fluorophores. We observe that ROS induce actin fragmentation and increase the number of polymerizing filaments. However, beyond a certain threshold, we see a transition from enhanced polymerization to the disassembly of actin networks. As such, via light, we have control over both actin assembly and disassembly in 3D.

## Localized severing leads to increase in actin polymerization

We use confocal fluorescence microscopy to observe the polymerization of fluorescently labeled actin in vitro. Our experimental setup is simple: we image a standard actin polymerization assay on a confocal microscope, where a square region of the sample, given by the microscope’s field of view, is exposed to light. Interestingly, prolonged exposure to the microscope’s light source causes an increase in filamentous actin (F-actin) in the illuminated region (Figure 1a), which we refer to as “actin printing”. We consistently observe light-induced F-actin accumulation within 10–15 minutes of continuous exposure across replicate experiments (for more details see Methods section). However, even a single, short, but intense exposure can result in an increase in actin polymerization if the sample is allowed to polymerize for several minutes before and after the light exposure (Figure S2a).

**Figure 1.**
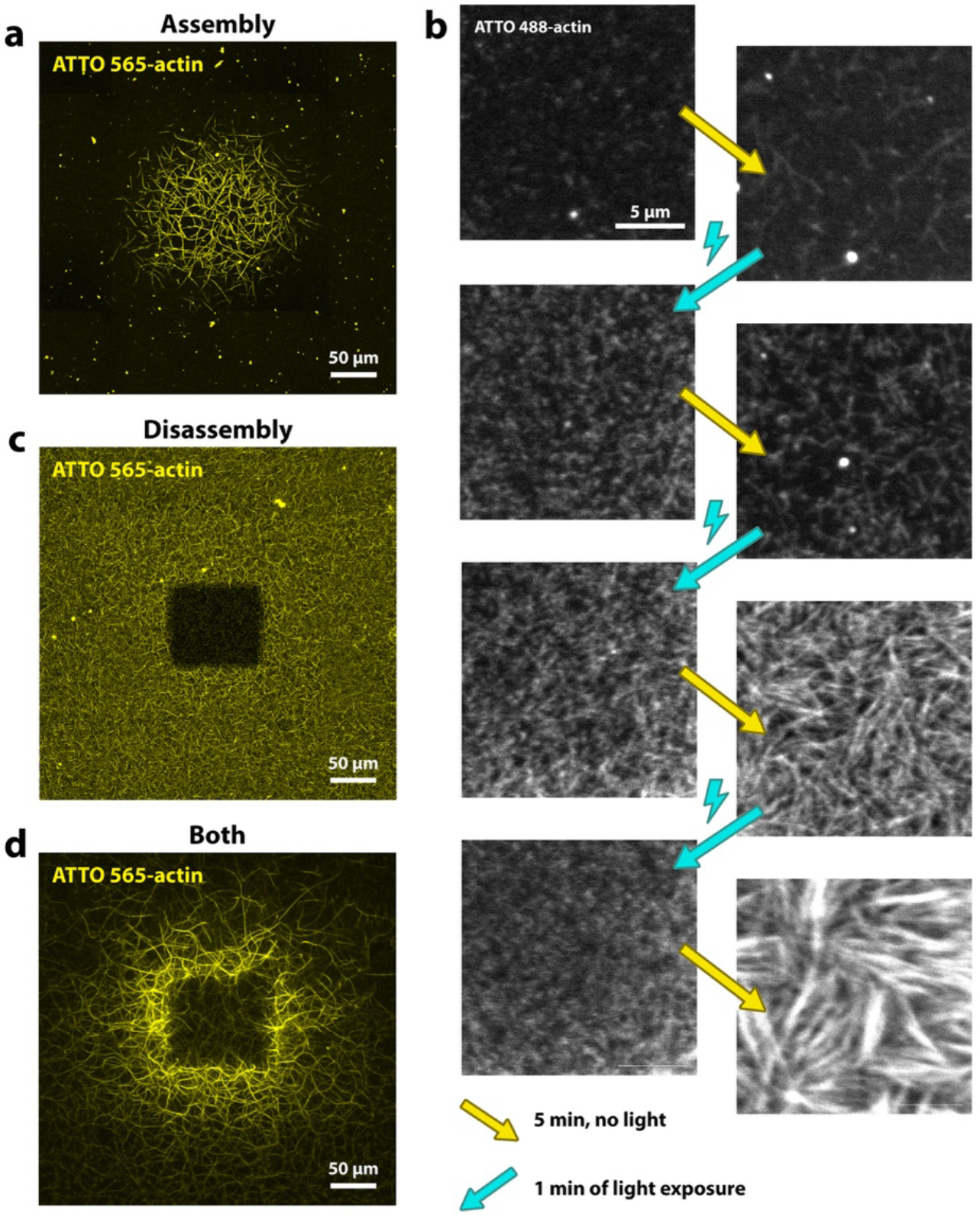
Light controlled actin assembly. **a)** Experiment showing light-induced increase in actin assembly. Condition with low intensity laser exposure and low actin concentration (1 µM). **b)** Intervals of high light exposure followed by rest time with no light exposure. Cyan arrows indicate strong light exposure, which causes filaments to fragment, resulting in numerous visible small speckles. The right column shows images after 5 minutes of no light exposure (yellow arrows) during which filament elongation occurs, which causes an increase in fluorescence intensity and structures that are visibly more filamentous. **c)** Actin dissembles in the light-exposed region in experiment with higher laser exposure, higher actin concentration (2 µM) and delayed onset of laser exposure. **d)** Combination of increased filament assembly and filament disassembly in experiment with high laser intensity and low actin concentration (1 µM). All images show maximum projections. In all experiments we include macromolecular crowder to induce actin bundling, even though this component is not required (Figure S1).

We gain first insights into the underlying mechanism by applying intervals of strong light illumination followed by periods without light: We observe that the light itself does not directly increase actin polymerization, but instead causes the fragmentation of existing filaments into smaller filaments. Only after subsequent dark periods, we observe an increase in F-actin, which exceeds that in areas not exposed to light (Figure 1b, Figure S2b).

## Excessive severing causes actin network disassembly

We can also achieve the opposite effect, in which light disassembles filaments in the exposed region. If we shine light on a sample with an existing network of F-actin, we see a slow disassembly of actin filaments in the immediate light-exposed area (Figure 1c).

Moreover, we also observe this effect in our actin printing experiments under polymerizing conditions, if we increase the light intensity. Above a certain light threshold we note a lack of filaments in the center of the final printed structure (Figure 1d). In these cases we initially observe an increase in actin polymerization in the exposed region, followed by network disassembly after several minutes (see Figure S3 and Movie 1). Filament disassembly typically does not extend far beyond the immediate light-exposed area. In contrast, light-induced increases in polymerization spread beyond this area, particularly when F-actin is disassembled in the illuminated area, which then results in the ring-shaped pattern as seen in Figure 1d.

The filament disassembly we observe resembles photo-bleaching, therefore we wanted to rule out that our observations might stem from such photochemical artifacts. Direct evidence comes from the brightfield images of our experiments, which can detect thick actin bundles and confirm the physical lack thereof in the center region of a ring-shaped print (Figure S4).

Time lapse recordings of exposure at high light intensities show a steady fragmentation of filaments into continuously smaller fragments (Movie S2), thus suggesting that filament severing is involved in both actin network assembly and disassembly.

## Fluorophore excitation results in actin printing

As described, with increasing light intensity we see a transition from filament assembly to filament disassembly in the light-exposed region. More precisely, we note a dependency of this behavior on the density of light, both over area and time (Figure S5a-c). Further, actin printing depends on the presence and concentration of fluorophores in our samples (Figure S5d-f). Our experiments require excitation wavelengths that match the fluorophore’s absorption spectrum (Figure S6), indicating that the excitation of fluorophores is critical part of the underlying mechanism.

In most of our experiments, we use fluorophores that are covalently bound to actin monomers, i.e. fluorescently labeled actin, which is commonly used to visualize actin for microscopy. Conveniently, in these experiments, these fluorophores double as a ‘photosensitizer’ for actin printing and as a microscopy label. We see both light-induced assembly and disassembly of F-actin for all different fluorescently labeled actins tested (Figure 2a and Figure S7).

**Figure 2.**
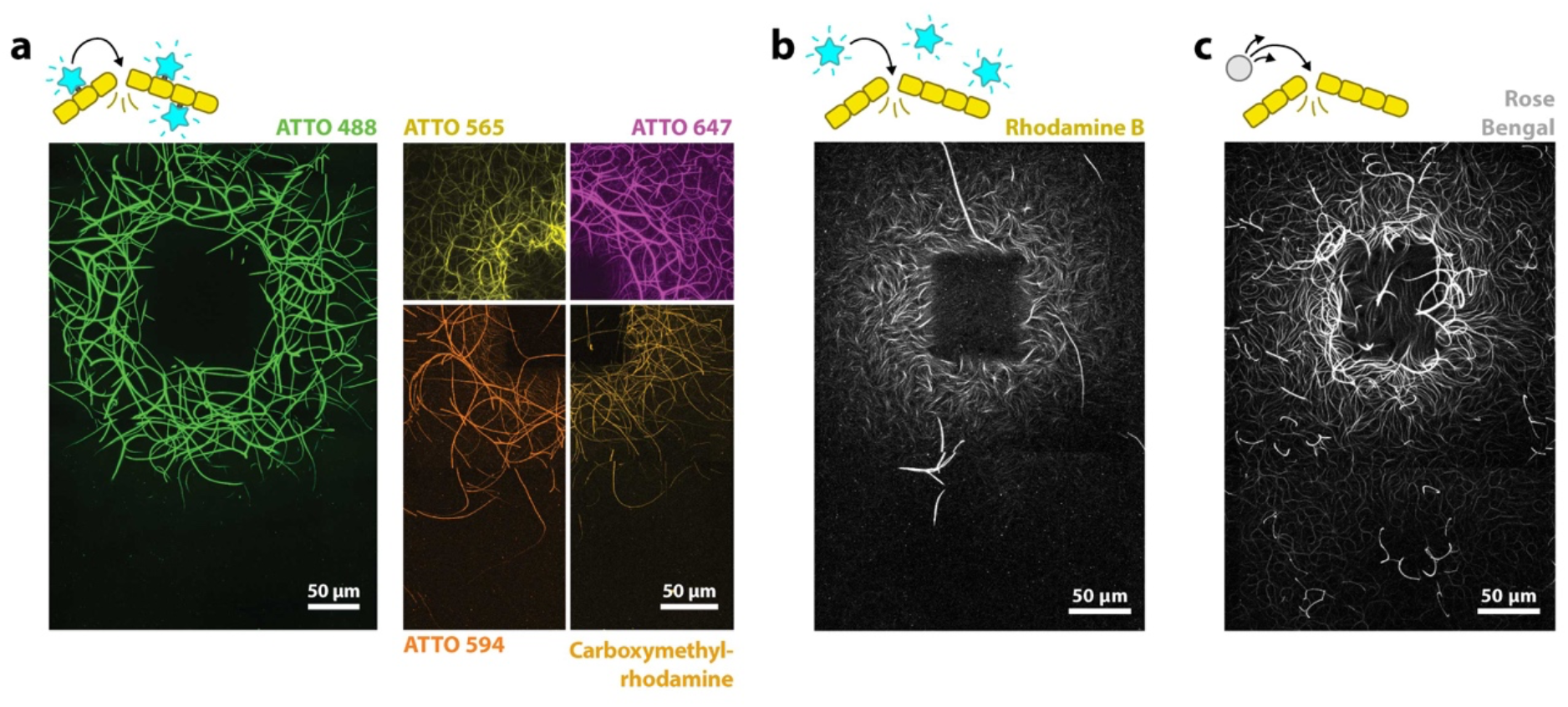
Various types of photosensitive molecules produce similar results. **a)** Ring-shaped prints are achieved with actin with 5 different fluorescent labels. These fluorophores are covalently bound to actin monomers. Various actin concentrations were used to achieve consistent outcomes (see Figure S7). **b)** Experiment with a fluorophore, which is not attached to actin (1µM actin, 12 µM rhodamine B). **c)** Similar to b), but instead of an efficient fluorophore, we use the photosensitizer rose bengal. Rose bengal produces ROS more efficiently and as such was used at much lower concentrations (1.5 µM actin, 0.04 µM rose bengal). In b) and c) we use low amounts of fluorescent labels for actin imaging, however with an excitation spectrum that does not overlap with the excitation wavelength used for printing. All figure panels show maximum projections of the samples.

While the use of fluorescently labeled actin simplifies the experiments, we note that fluorophores do not have to be covalently bound to actin molecules: We see both increases in polymerization as well light-induced disassembly when we instead have free fluorophores in solution, which are not bound to actin. Figure 2b shows an experiment with unbound rhodamine B. However, here we have to increase the concentration of fluorophores to achieve results consistent with those with fluorescently labeled actin, indicating that proximity of the fluorophores to the filaments affects the outcome.

Instead of an efficient fluorophore, we can use rose bengal, a compound used as a photosensitizer, which produces high amounts of ROS when excited with light. We observe rose bengal to be particularly potent in our experiments. To achieve light-induced assembly and disassembly, only a fraction of the concentration of rose bengal is necessary compared to fluorophores like rhodamine B (Figure 2c). In experiments in which we do not decrease the rose bengal concentration, we observe an extreme degree of actin disassembly that spreads over an area with a radius of over 1 mm (Figure S8).

Actin polymerization is temperature dependent, therefore we test whether our actin printing effect is caused by a local increase in temperature. We replace fluorophores with gold nanoparticles, which produce thermal energy when excited within their absorption range.^15^ In these experiments we do not observe actin printing effects, indicating that the effect is not based on increases in temperature (Figure S9).

It is widely understood that fluorophores produce ROS upon excitation with light,^16^ which in turn can oxidize biomolecules such as proteins.^17^ Oxidation has damaging effects, often inducing conformational changes that interfere with protein function. As such, ROS might react with and damage actin monomers within filaments, which could explain our observation of filament severing. To test whether actin printing relies on ROS, we include antioxidants and oxygen scavengers in our samples. We test 3 different reducing agents (β-mercaptoethanol, p-phenylenediamine and FluMaXx, a commercial enzymatic oxygen scavenger system) which either prevent ROS formation or neutralize formed ROS. We find that all three can prohibit actin printing (Figure S10).

## Modeling explains how severing drives both assembly and disassembly

Based on our observations, we propose that the fluorophores in our experiments generate ROS upon excitation with light. ROS lead to the severing of actin filaments, most likely due to oxidation of actin monomers within filaments. Given the nucleation-limited nature of actin polymerization, this would explain an increase in F-actin: Filament severing results in an overall increase in filaments, and therefore in a greater number of polymerizing ends. Because continued polymerization of existing filaments is kinetically favored over the nucleation of new filaments, an increase in filaments directly correlates with an increase in actin polymerization. The schematic in Figure 3a illustrates this proposed mechanism. Severing also offers an explanation for network disassembly: If ROS concentrations are high enough, filament disassembly dominates over polymerization and F-actin dissipates in the light-exposed area.

**Figure 3.**
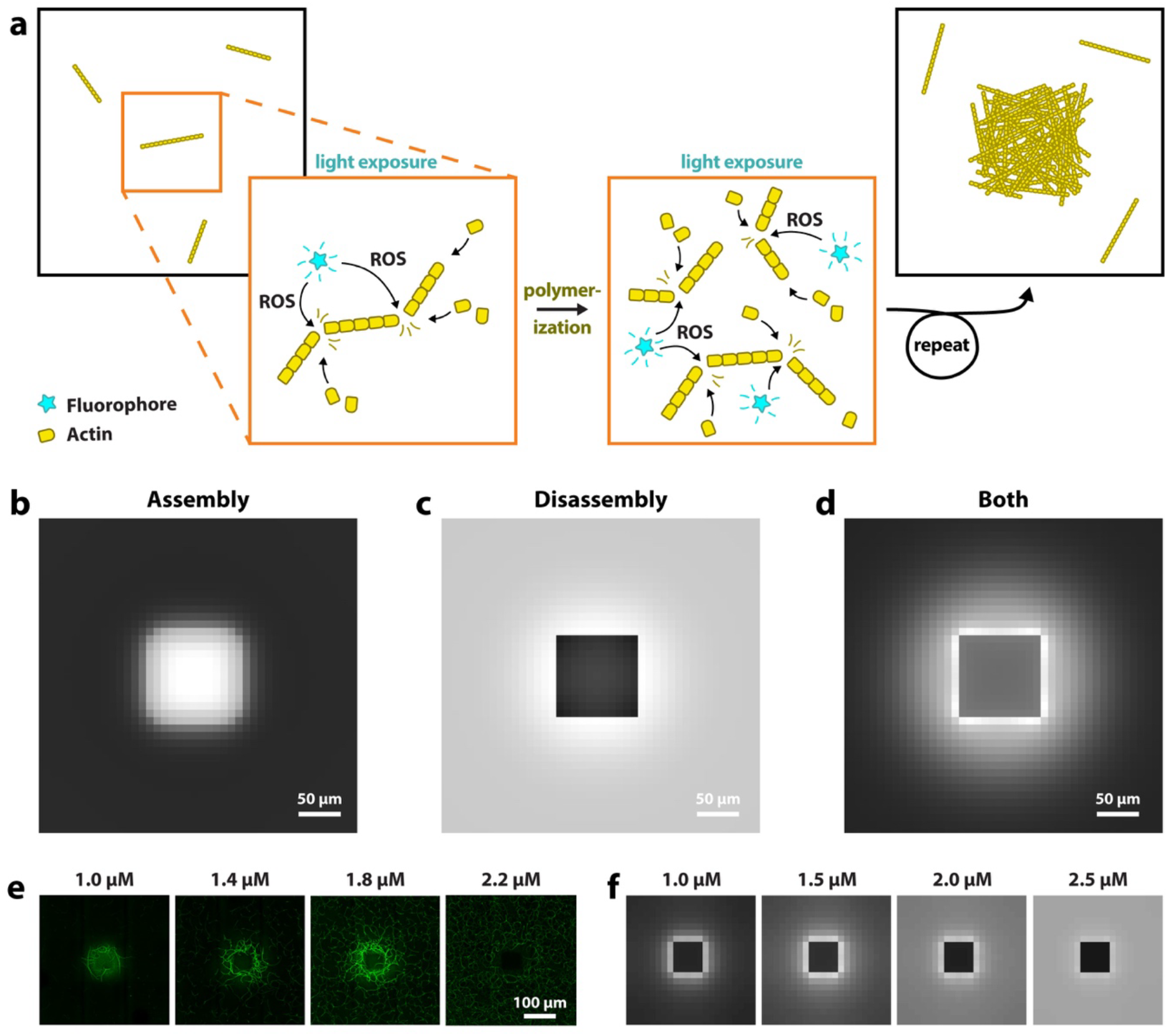
Model based on actin severing. **a)** Schematic illustrating the increase in actin assembly through light- induced severing via reactive oxygen species (ROS). A region within a polymerizing actin sample is exposed to light. Fluorophores produce ROS, which oxidize actin molecules within filaments and thus cause filament fragmentation. Oxidized monomers likely dissociate (not shown) and polymerization continues. Actin assembly is nucleation-limited, thus an increase in the number of filaments increases polymerization. **b)**, **c)** and **d)** show results of our computational model. Simulations correspond to experiments shown in Figure 1 a), c) and d). **e)** Experiments with Atto488-labeled actin at different actin concentrations. **f)** Simulations show a similar concentration-dependent behavior.

To test the hypothesis that ROS-mediated filament severing underlies both enhanced filament assembly as well as their disassembly, we developed a continuum 2D mathematical model that incorporates actin nucleation and growth. Assuming that local filament severing rate is proportional to local ROS production rate within the illumination region, we evolve the concentration distributions of monomers and filaments, accounting for their length-dependent diffusion. See methods for more details.

In agreement with our experiments, at moderate light intensity, the model shows that fragmentation of actin filaments increases the number of polymerizing filaments and therefore results in an increase in F-actin concentrations in the light-exposed area (Figure 3b and supplementary Figure S11). In further agreement with experiments, light exposure can also lead to an overall decrease in F-actin, because continued fragmentation leads to filaments that are small enough to diffuse out of the illuminated area (Figure 3c). If under polymerizing conditions, once these small filaments reach the periphery, they continue to polymerize and become too long to diffuse back into the exposed region, which explains the ring-shaped pattern of increased F-actin (Figure 3d).

Our simulations give us further insights into whether other, additional contributors play a role in the light-induced actin network disassembly. Several factors could cause a lack of monomeric actin (G-actin) in the light-exposed area, therefore limiting actin polymerization. In the light-exposed area, G-actin is quickly used up and the diffusion of new G-actin might be rate-limiting. To test this, we speed up simulated G-actin diffusion, but only find this to cause significant differences for very large areas of exposure (Figure S12). Next, we tested whether oxidative damage of actin could reduce the pool of intact G-actin. We note that in our simulations, the pool of oxidized actin stays very small compared to that of intact actin monomers (Figure S13), suggesting that protein damage does not cause a significant decrease in actin polymerization.

We investigate the concentration dependence of actin printing and compare our model to experimental results. Both experiments and simulations show a similar printing behavior across a range of actin concentrations, with less light-induced increases in actin polymerization at higher actin concentrations (Figure 3d). Actin nucleation follows a power law relationship with actin concentration; as such, at higher concentrations a majority of the actin is already polymerized at the onset of light exposure.

## Printing complex cytoskeletal networks in 3D

We find the printing effect to be scalable and to allow for patterns in various shapes and sizes. In Figure 4a and b we show printed structures between ∼ 1.6 mm and ∼ 60 µm in width. Figure 4a and Figure S14 show large actin prints achieved with a widefield microscope.

**Figure 4.**
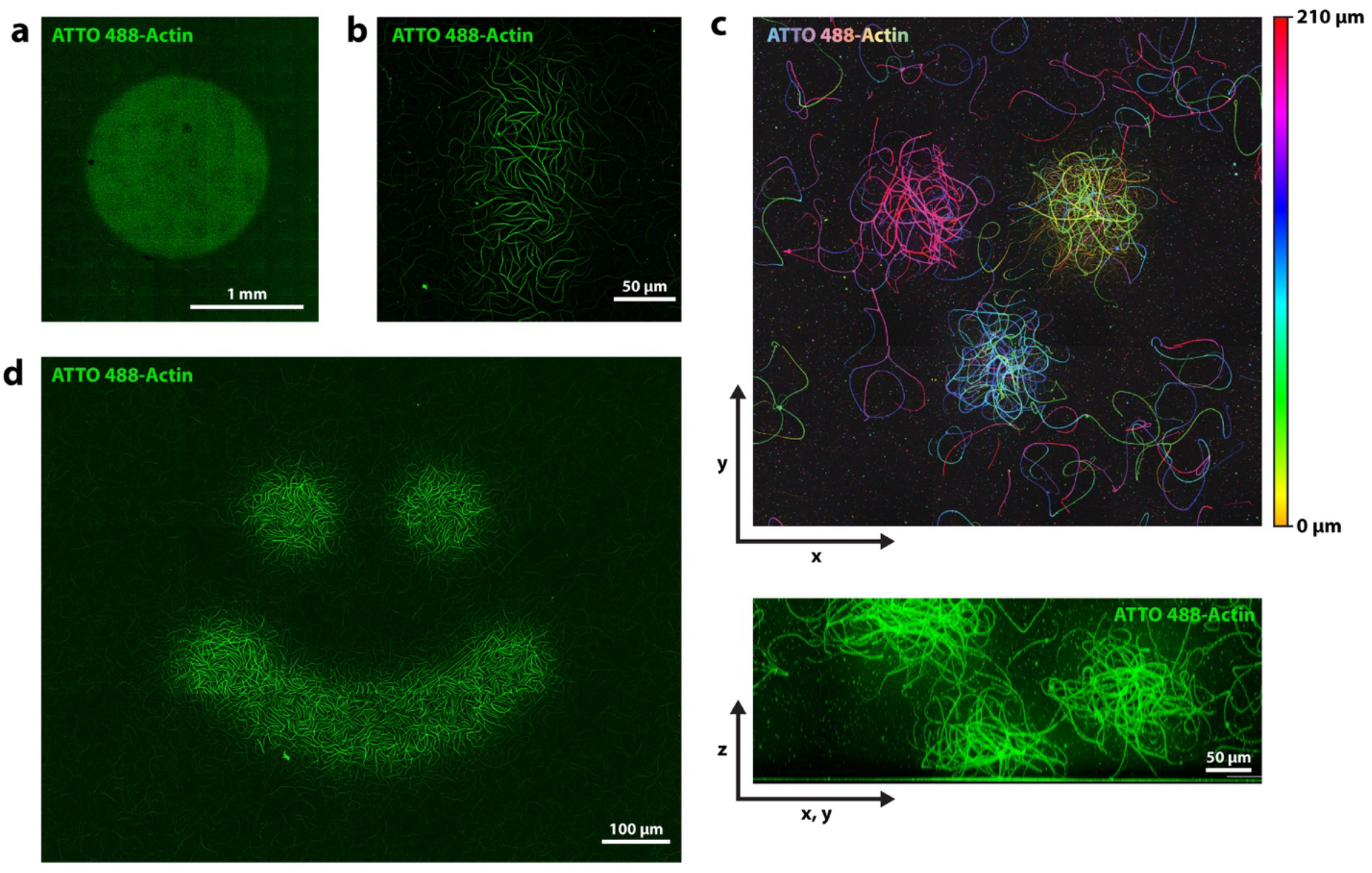
Spatial control over actin assembly. **a)** Light exposure via illumination with a widefield microscope. A large circular area of ∼1.6 mm in diameter was exposed, leading to increased actin assembly within the area. **b)** Light exposure via confocal laser scanning microscope of a small region of 60 µm x 140 µm in size. **c)** Confocal microscopes allow for control of actin polymerization in 3D. Shown are three actin patches polymerized by focusing the confocal beam to 3 different z-heights. Top image: false color image, color represents height in z. Bottom image: side view (projected diagonally). **d)** Arbitrary patterns such as this smiley shape can be printed. All experiments use 1.5 µM actin.

Our technique allows for control over actin assembly in 3D: Using the confocal microscope, we can expose the sample at different positions in z, which results in patches of actin being printed at different z-heights. Figure 4c shows a false color image indicating the position in z, of an experiment where we exposed the sample in 3 independent locations at 3 different z-heights.

Rather than limiting illumination to simple squares or circles, we can project more complex patterns onto the sample. As an example, Figure 4d shows the sample illuminated in a “smiley” pattern.

We find that not only eukaryotic actin is affected by ROS-induced severing, but also the bacterial actin homolog ParM. By combining eukaryotic actin and ParM, we can achieve what we call multi-color printing. We combine ATTO 565-labeled actin and Alexa Fluor 488-labeled ParM and exposed the sample with both excitation lasers at high intensities, which results in an alternating pattern, with ParM in the exposed area, surrounded by an actin ring-print, followed by ParM further away (Figure 5). As Figure 5b confirms, we observe a type of anti-correlation, where the concentrations of one type of polymer are low in regions dominated by the other (also see Figure S15). While we show one specific case here, different variations of this experiment are possible. In supplementary Figure S16 we highlight more observations from these experiments with both biopolymers. Movie 3 and supplementary Figure S17 show the time course within the exposed region of such an experiment, which also unveils some interesting aspects.

**Figure 5.**
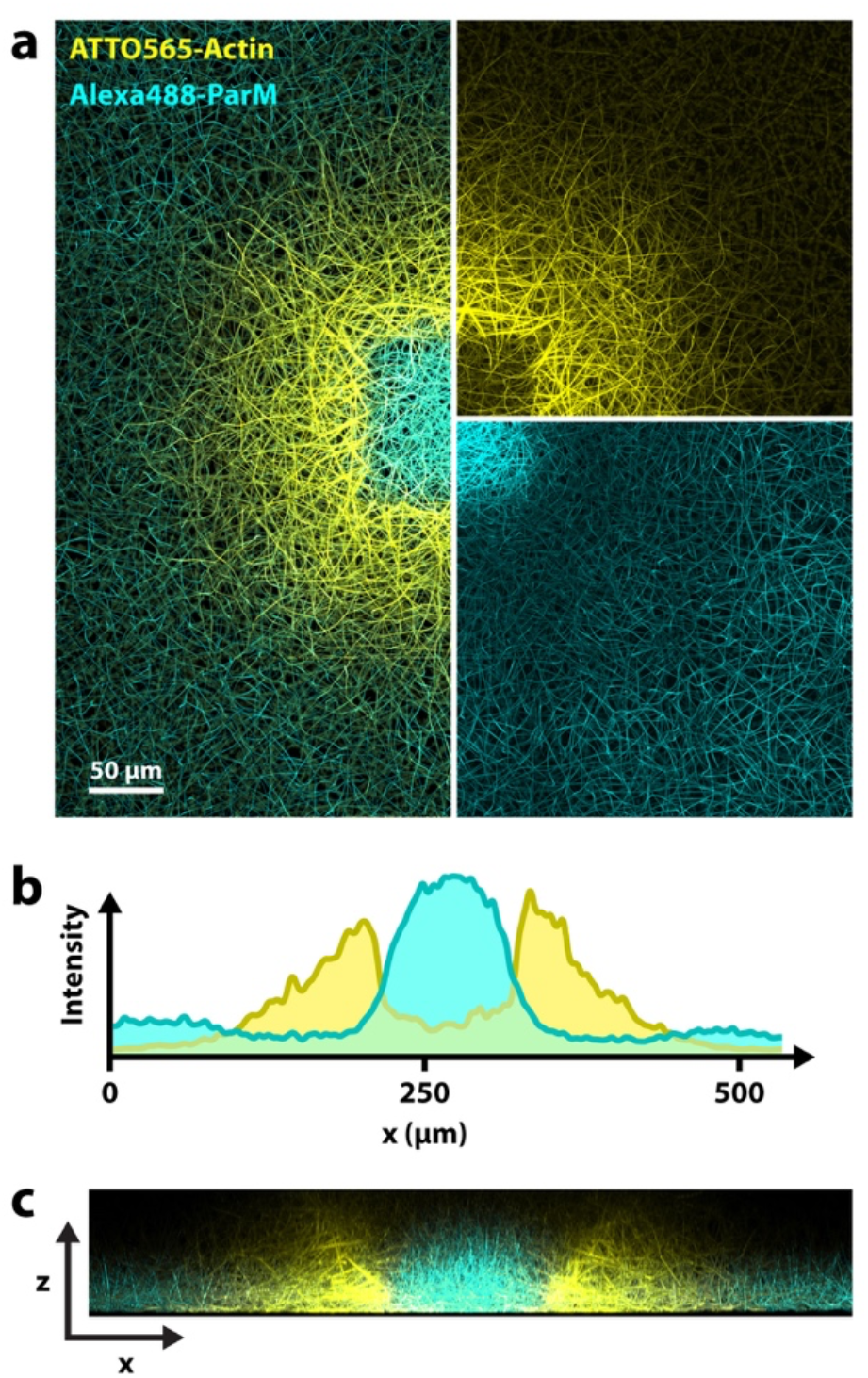
Multi-color printing. Experiment with two types of actin: Eukaryotic actin and the bacterial actin homolog ParM. (Maximum projection.) Both are affected by laser excitation and can be printed. In this case, ParM polymerization was enhances in the center and actin polymerization in the periphery (ring-shaped print). **a)** x-y-view with both channels combined on the left and split channels on the right (top: actin, bottom: ParM). **b)** Fluorescence intensity profile measured horizontally across the center. **c)** Side view (x,z) cross section of the sample. (Maximum projection of the light-exposed region.) Different morphologies of the two different cytoskeletal networks are visible.

## Discussion

We demonstrate that via the excitation of fluorophores we can control the assembly and disassembly of the cytoskeletal protein actin in three dimensions, including the capability for ‘multi-color’ printing. Our experimental data and simulations support a mechanism in which ROS are produced by these photosensitive molecules, which then fragment actin filaments.

We think our findings are significant for various fields. The ability to 3D print actin hydrogels introduces new possibilities for biomaterials science, while the underlying mechanism of growth via fragmentation could be applicable to other materials. Further, our results are a consideration for experiments involving fluorescently labeled actin, where this phenomenon could lead to artifactual effects. Moreover, controlling actin network geometry is a valuable tool for in vitro reconstitution studies,^18^ where thus far, spatial control of actin was limited to 2D surface patterning techniques.^19-21^ Perhaps most importantly, we believe this ROS-dependent mechanism likely affects actin remodeling in living cells, where all requirements are met for it to take place.

In previous literature, we find reports that are consistent with our findings. In vitro, fluorescently labeled actin exposed to light has been observed to cause an increase in actin polymerization, which previously was either attributed to the light^22^ or the fluorescent label alone^23,24^ but not the combination thereof. On the other hand, high light intensities were associated with ‘photodamage’ and a collapse of actin networks in vitro.^25-29^

In vivo, correlations between ROS and actin polymerization are well known and the importance of redox regulation of actin has become of interest.^10-12,30,31^ Several studies show that ROS are central for actin remodeling in cells: depleting cells of ROS leads to actin cytoskeleton disassembly and impedes actin-dependent functions such as cell migration,^32,33^ wound healing^34^ and neuronal growth^35,36^. Supplementation with external ROS can rescue this actin and cell function loss.^37^ Bulk exposures to ROS have also been employed in earlier studies on oxidative stress and are described lead to an increase in F-actin in various cell types like macrophages^40-43^, astrocytes,^44^ and drosophila embryonic cells^45^. More localized generation of ROS has been achieved by tagging actin with an engineered ROS-generator, which results in a drastic increase in F-actin in various types of cells.^38,39^ Further, a strong correlation has been observed between ROS generation and actin network density around mitochondria: Damaged mitochondria produce high levels of ROS, leading to ‘acute damage-induced actin’ ^46,47^ and certain environmental changes have been shown to correlate with both increases in mitochondrial ROS and peripheral actin^48^.

These studies link increases in ROS to increases in actin, yet none offer conclusive explanations for this connection. On the other hand, there are numerous reports of a negative correlation between ROS and actin, which typically are credited to the general destructive nature of ROS. Excitingly, both actin assembly and disassembly can be explained by our model via ROS-induced severing. We assume fragmentation can both enhance polymerization as well as depolymerization, depending on which process is favored by the current environment. A second effect is that fragmentation increases filament diffusion, which explains local disassembly in our experiments, even under polymerizing conditions. In cells, the consequences of ROS-induced fragmentation are likely dependent on many different factors, including the availability of actin monomers and specific actin regulators. A relevant comparison is the actin-severing protein cofilin, which has been shown to increase actin polymerization^49-51^ but usually collaborates with other proteins to promote filament disassembly.^52^ Thus, we speculate that ROS-induced severing might have extensive but complex implications for actin remodeling in cells. And indeed, many recent studies describe the effects of ROS on actin rather broadly, i.e. that it affects actin remodeling, reorganization or increases actin dynamics across various cellular processes,^12,53^ such as cell migration,^54-56^ cell adhesion^57-61^ and cell growth^62^.

Much of actin research has focused on the numerous actin accessory proteins that specifically bind and regulate actin polymerization and network architecture, offering an explanation for the remarkable versatility of the actin cytoskeleton. Our findings broaden the understanding of actin organization beyond specific protein interactions, suggesting that ROS, which are ubiquitous in cells and integral to cell homeostasis, add another layer of regulation to actin dynamics and remodeling.

## Methods

### Proteins

All eukaryotic proteins, namely actin (alpha-actin skeletal muscle, rabbit), including labeled actins (alpha-actin skeletal muscle, rabbit) and fascin (human, recombinant) were purchased from HYPERMOL (www.hypermol.com). ParM was provided by the Sourjik lab and purified according to Hürtgen et al., 2019.^63^ In short, BL21(DE3) cells were transformed with pET11a protein expression vectors under the control of a T7 promoter. Cells were lysed by sonication, clarified by centrifugation and subsequently subjected to an ammonium sulfate cut. After further centrifugation the lysate was treated with ATP solution to induce polymerization. These polymers were centrifuged again, resuspended and gel-filtered through a Superdex S200 column. Pure fractions were then determined by SDS-PAGE, pooled, and frozen at −80 °C.

### Reaction mix preparation

For a typical experiment, a 23 µl reaction mix was prepared. First, 9 µl of a concentrated buffer solution containing all components except protein was pipetted into a microcentrifuge tube. Milli-Q ultrapure water (Millipore) was then added to adjust the volume such that the final total volume - including subsequent additions - would be 23 µl. Next, 0.5 µl of 100 mM ATP or ATP-γ-S was added, followed by the desired volume of actin from a 23 µM stock solution (1 mg/ml); for example, 1.5 µl was added to achieve 1.5 µM of actin. Typically we use 70% labeled actin, but lower fractions are possible (Figure S5). A typical final composition of the reaction mix is: 26 mM Tris-HCl (pH 7.5), 0.9 mM DTT, 0.01 mM CaCl_2_, 87.0 mM KCl, 1.74 mM MgCl_2_, 0.43% methyl cellulose, 2.2 mM ATP-γ-S, and 1.5 µM actin. In most of our experiments we use ATP-γ-S, however observe no difference when compared to ATP (see Figure S19). Actin concentrations vary between experiments, as we note in respective figure captions.

The experiments with ParM (Figure 5, Figure S15-S19, Movie 3), contain 2 µM actin (70% labeled), 22 µM ParM (35% labeled) and 15 mg/ml bovine serum albumin in addition to the above mentioned reaction mix components.

### Slide preparation

Slide chambers were assembled as illustrated in Supplementary Figure S18. Unless otherwise noted, experiments were conducted using a 22 × 32 mm #1.0 cover glass (Kemtech, Microscope Cover Glass) as the base. An adhesive spacer with circular cut-out (Grace Bio-Labs, SecureSeal™ Imaging Spacers, 9mm DIA x 0.12mm Depth, 18 × 18MM OD) was then applied to form the reaction chamber. A 9 µl volume of reaction mix was pipetted into the chamber and sealed with a 22 × 22 mm cover glass. The pipetted volume was chosen to slightly exceed the chamber volume to ensure complete filling. It is best to avoid air bubbles in the chamber or generally an asymmetric setup, as this can create directional flow during exposure/imaging and thus can lead to an asymmetric actin print. For large-scale prints (Figure 4a and Supplementary Figure S14a), the chamber height was increased by stacking multiple adhesive spacers.

### Light Exposure

We use a Leica SP5 confocal laser scanning microscope for both light exposure and image acquisition for all experiments with exception of the following: Experiments in Figure 1b, S8b and Movie XY were performed on a ZEISS LSM 800 confocal laser scanning microscope; experiments in Figure 5, S15-S17 and S18b were performed on a ZEISS LSM 780 confocal laser scanning microscope; Experiments in Figure 4a and Figure S14 were performed on a Leica Thunder widefield microscope.

We typically incubate the reaction mix for 10-20 minutes before light exposure, to allow a small amount of actin to polymerize before the light-induced severing, which initially accelerates actin printing. For most additive actin prints, we expose the sample for 60 minutes at 0.2 mW laser power (roughly 3% laser power on a modern confocal microscope) with 50% on-time using the scanning laser of the confocal microscope. However, for larger actin prints, we decrease the light exposure, by decreasing on-time, and extend exposure time. For Figure 4a, Figure 4d and Figure S14a we performed 5 hour, 3 hour and 10 hour long exposures respectively. For subtractive and ring-shaped prints we exposure the sample for 90 minutes or more at 0.2 mW laser power with 50% on-time using the scanning laser of the confocal microscope. These experiments are not sensitive to over-exposure, i.e. a longer exposure to light will not alter the result significantly.

### Model

To interpret our experimental results, we implemented a reaction–diffusion model based on partial differential equations. The system is modeled in 2D, assuming uniformity along the optical axis due to the thin sample geometry. Reflecting boundary conditions are applied at the edges of the simulation domain. We track spatial and temporal concentrations of non-oxidized monomers *A*(**r**, *t*), oxidized monomers *O*(**r**, *t*), and filaments of length *N* ≥3, denoted Ψ_*N*_(**r**, *t*). Filaments nucleate as trimers with rate *k*_*nuc*_,^64^ grow via monomer addition with rate *k*_*pol*_, and diffuse with an effective diffusion coefficient *D*_*N*_ that decreases with filament length.^65^ Oxidative severing occurs locally in the laser-exposed region and releases oxidized monomers. To prevent excessive filament growth, we also include spontaneous fragmentation with rate *k*_*frag*_.^66-68^

For filaments longer than trimers, the evolution equation is:

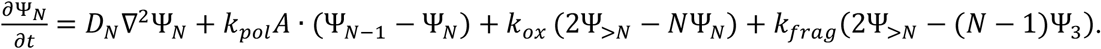

The terms correspond to filament diffusion, elongation, severing by oxidation, and spontaneous fragmentation, respectively. The oxidation rate *k*_*ox*_ is proportional to total actin concentration and only nonzero within the illuminated region during laser exposure.

This model captures the essential features of actin network dynamics under oxidative severing within a square region. A complete description, including initial conditions, numerical methods, and a table with parameter values, is provided in the supplements.

## Acknowledgements

We thank Nadab Wubshet, Davide Bray and Jingcheng Tan for help with experiments. We thank David Rutkowski for help with computational methods. We thank the lab of Victor Sourjik, including Judita Mascarenhas and Daniel Hürtgen, for providing us with purified ParM protein. We thank the following people for helpful discussions: Andreas Manz, Arnaud Echard, Charlotte Aumeier, Christine Field, Daniel Needleman, Daniel Oblinsky, David Kast, Guillaume Romet-Lemonne, Gregory Scholes, Petra Schwille, Ricardo Rizzo, Sylvia Jansen and Tim Mitchison. DV was supported by NIH grant R35GM136372 and a Lehigh Physics Faculty Research Innovation Fellowship.

