## Supplements for "3D printing cytoskeletal networks: ROS-induced filament severing leads to surge in actin polymerization"

### Supplementary Information

#### Model

*Setup and initialization.* We use a partial differential equations model to describe the evolution of the concentration of non-oxidized actin monomers $A(\mathbf{r},t)$, oxidized actin monomers, $O(\mathbf{r},t)$, actin filaments with $N \geq 3$ subunits, $\Psi_N(\mathbf{r},t)$, and barbed ends (or, equivalently, total number of filaments), $B(\mathbf{r},t) = \sum_{N=3}^{\infty} \Psi_N(\mathbf{r},t)$, at position $\mathbf{r}$ and time *t*. A useful quantity in the model is the concentration of filaments with more than $N$ subunits: $\Psi_{>N}(\mathbf{r},t) = \sum_{N'=N+1}^{\infty} \Psi_{N'}(\mathbf{r},t)$. We use a 2D approximation, considering that the experimental system behaves approximately uniformly along the direction normal to the confining glass slides. Reflecting boundary conditions are applied to the outer boundaries of the simulation box. We initialize the system such that $A(\mathbf{r},0) = A_0$ is equal to the initial actin monomer concentration, with all other concentrations set to 0. The model parameter values are shown in Table 1.

*Laser Intensity and Oxidation.* We assume the laser is applied uniformly within a square region of side length $2a$, following an initial incubation period of duration $t_{inc}$. Radicals causing actin oxidation are assumed to be generated locally, in proportion to the total local actin concentration. Assuming the process of radical creation, diffusion, and oxidation occurs at sub-μm distances,[1] smaller than the resolution of our space discretization, the local rate of actin oxidation is: $k_{ox}(\mathbf{r},t) = k_{ox}^{0}[A(\mathbf{r},t) + F(\mathbf{r},t) + O(\mathbf{r},t)]$ for $|x|, |y| < a$ , $t < t_{inc}$, and zero otherwise. Here $k_{ox}^{0}$ is a constant proportional to the laser intensity and $F(\mathbf{r},t) = \sum_{N=3}^{\infty} N\,\Psi_N(\mathbf{r},t)$ is the total F-actin concentration.

*Nucleation.* New filaments are assumed to nucleate as trimers with a rate constant $k_{nuc}$. This assumption can empirically fit the kinetics of actin polymerization over a sufficiently narrow range of concentrations.[2] Here we don't attempt to describe the precise pathway of nucleus formation[3] or role of ATP hydrolysis,[4] and we use trimers rather than tetramers (which may provide a more accurate concentration dependence[2] to simplify the equation below.

*Polymerization.* Filaments are assumed to elongate with rate constant $k_{pol}$. We neglect the effects of depolymerization from either end; thus, at long times evolves towards $A = 0$ instead of evolving towards a critical concentration much smaller than the initial $A_0$.

*Severing and fragmentation.* Oxidation of an actin subunit in the middle of a filament is assumed to result in severing of the actin filament, release of the oxidized subunit into the oxidized monomer pool, and creation of a new barbed and pointed end. When oxidation happens at the third subunit from either end, we assumed this results in the release of one oxidized and two non-oxidized monomers. When oxidation happens at the penultimate subunit from either end, we assume this results in the release of one oxidized and one non-oxidized monomer. When oxidation happens at a terminal subunit, this subunit is released as oxidized monomer. While these specific assumptions are not essential to the results, they need to be postulated for mass conservation in the simulations. In addition to severing by oxidation, we also implement spontaneous filament fragmentation with rate constant $k_{frag}$ between actin subunits. Spontaneous fragmentation is needed for accurate description of actin polymerization kinetics[5-7] and restricts the formation of extremely long filaments in the absence of oxidative severing. Fragmentation near the ends is treated similar to oxidative severing, but without the release of oxidized monomer.

*Diffusion.* We assumed that translational filament diffusion decreases linearly with length,[8] up to a maximum length, above which the diffusion can be neglected due to confinement by methylcellulose or bundling with nearby actin:

$$D_N = \begin{cases} 2D_A/N, & N < N_{freeze} \\ 0, & N \geq N_{freeze} \end{cases} . \quad (1)$$

Here $D_A$ in the diffusion coefficient of actin monomers and the factor of 2 is for the two strands of actin filaments.

*Model equations.* The monomer concentration changes over time, according the following equation that accounts for monomer diffusion, nucleation, polymerization, and release of monomers from filaments after oxidative severing or spontaneous fragmentation:

$$\frac{\partial A}{\partial t} = D_A \nabla^2 A - 3\, k_{nuc} A^3 - k_{pol} A \cdot B - k_{ox}\, A + \; r_{sever} + r_{frag}\, . \quad (2)$$

Here $r_{sever}(\mathbf{r}, t)$ and $r_{frag}(\mathbf{r}, t)$ are the rates of actin monomer release after oxidative severing or spontaneous fragmentation at the third, penultimate, or terminal subunits of long and short filaments:

$$r_{sever} = k_{ox}(4\Psi_{>5} + 2\Psi_{>4} + 4\,\Psi_5 + 6\Psi_4 + 6\Psi_3)$$
$$r_{frag} = k_{frag}(4\Psi_{>4} + 2\Psi_{>3} + 4\,\Psi_4 + 6\Psi_3). \quad (3)$$

The differences in the prefactors of the oxidative severing and fragmentation terms are due to counting one less fragmentation interface compared to number of monomers. For trimers, we have:

$$\frac{\partial \Psi_3}{\partial t} = D_3 \nabla^2 \Psi_3 + 3\, k_{nuc} A^3 - k_{pol} A \cdot \Psi_3 + k_{ox}\,(2\Psi_{>3} - 3\Psi_3) + k_{frag}(2\Psi_{>3} - 2\Psi_3)\, . \quad (4)$$

For filaments longer than trimers:

$$\frac{\partial \Psi_N}{\partial t} = D_N \nabla^2 \Psi_N + k_{pol} A \cdot (\Psi_{N-1} - \Psi_N) + k_{ox}\,(2\Psi_{>N} - N\Psi_N) + k_{frag}(2\Psi_{>N} - (N-1)\Psi_3). \quad (5)$$

Finally, the concentration of oxidized actin, which is assumed to accumulate, being unable to polymerize, obeys:

$$\frac{\partial O}{\partial t} = D_A \nabla^2 O + k_{ox}\,(A + F) \quad (6)$$

*Testing the role of actin monomer diffusion.* To examine the role of monomeric actin diffusion limitation in establishing negative printing, we evolved the model as above, but we homogenized the actin monomer concentration field, after each time step of the simulation, by replacing $A(\mathbf{r}, t)$ by its average value in space.

*Numerical Solution.* We evolved the concentrations $\Psi_N$ up to an $N$ of order 15,000, checking that this number was large enough to not influence the results of the simulations.

Reaching this upper length limit was assumed to stop polymerization. We found that a simple Euler scheme on a square lattice of unit size $dx$ gave a stable solution, for time step of order $dt = 0.05$ s. We checked that the total mass in the system was conserved with accuracy better than 1% over the course of a simulated experiment. We note the approximation that the model evolves the local concentration of filaments, even if their length may exceed the unit lattice size.

| Parameter | Value | Justification |
|---|---|---|
| $D_A$ $(\mu m^2 s^{-1})$ | 50 | G-actin diffusion coefficient, close to value for water,[9] assuming weak influence of methylcellulose. |
| $k_{pol}$ $(\mu M^{-1} s^{-1})$ | 10 | Typical value for actin polymerization in vitro |
| $k_{nuc}$ $(\mu M^{-2} s^{-1})$ | $8 \cdot 10^{-8}$ | Slightly larger than $8 \cdot 10^{-9}$ used for actin polymerization kinetics in bulk pyrene assays.[7] This larger value needed to capture the range of actin concentrations in this study, may reflect the different ionic conditions, presence of methylcellulose, and confinement by the glass slide. |
| $k_{frag}$ $(s^{-1})$ | $5 \cdot 10^{-8}$ | Similar to value used to fit bulk pyrene assays.[7] |
| $k_{ox}^0$ $(\mu M^{-1} s^{-1})$ | $10^{-5} - 10^{-3}$ | Varied to capture the range of positive and negative printing in this work as function of laser intensity |
| $t_{inc}(s)$ | 720 | Typical incubation period prior to laser exposure in this work |
| $N_{freeze}$ (mon.) | 187 | Estimated |
| Box size (μm) | 240 | |
| $a$ (Laser box half size, μm) | 24 | |
| $dx$ (Lattice size, μm) | 6 | |

**Table 1**: Model Parameter Values


1 Mattila, H., Khorobrykh, S., Havurinne, V. & Tyystjärvi, E. Reactive oxygen species: Reactions and detection from photosynthetic tissues. *Journal of Photochemistry and Photobiology B: Biology* **152**, 176-214 (2015). https://doi.org/https://doi.org/10.1016/j.jphotobiol.2015.10.001

2 Tobacman, L. S. & Korn, E. D. The kinetics of actin nucleation and polymerization. *J Biol Chem* **258**, 3207-3214 (1983).

3 Sept, D. & McCammon, J. A. Thermodynamics and Kinetics of Actin Filament Nucleation. *Biophysical Journal* **81**, 667-674 (2001). https://doi.org/10.1016/S0006-3495(01)75731-1

4 Rosenbloom, A. D., Kovar, E. W., Kovar, D. R., Loew, L. M. & Pollard, T. D. Mechanism of actin filament nucleation. *Biophysical Journal* **120**, 4399-4417 (2021). https://doi.org/10.1016/j.bpj.2021.09.006

5 Schmoller, Kurt M., Niedermayer, T., Zensen, C., Wurm, C. & Bausch, Andreas R. Fragmentation Is Crucial for the Steady-State Dynamics of Actin Filaments. *Biophysical Journal* **101**, 803-808 (2011). https://doi.org/10.1016/j.bpj.2011.07.009

6 Sept, D., Xu, J., Pollard, T. D. & Andrew McCammon, J. Annealing Accounts for the Length of Actin Filaments Formed by Spontaneous Polymerization. *Biophysical Journal* **77**, 2911-2919 (1999). https://doi.org/10.1016/S0006-3495(99)77124-9

7 Heisler, D. B. *et al.* ACD toxin–produced actin oligomers poison formin-controlled actin polymerization. *Science* **349**, 535-539 (2015). https://doi.org/doi:10.1126/science.aab4090

8 Doi, M. & Edwards, S. F. *The theory of polymer dynamics*. New edn, (Clarendon, 1986).

9 Tait, J. F. & Frieden, C. Polymerization and gelation of actin studied by fluorescence photobleaching recovery. *Biochemistry* **21**, 3666-3674 (1982). https://doi.org/10.1021/bi00258a022

### Supplementary Figures

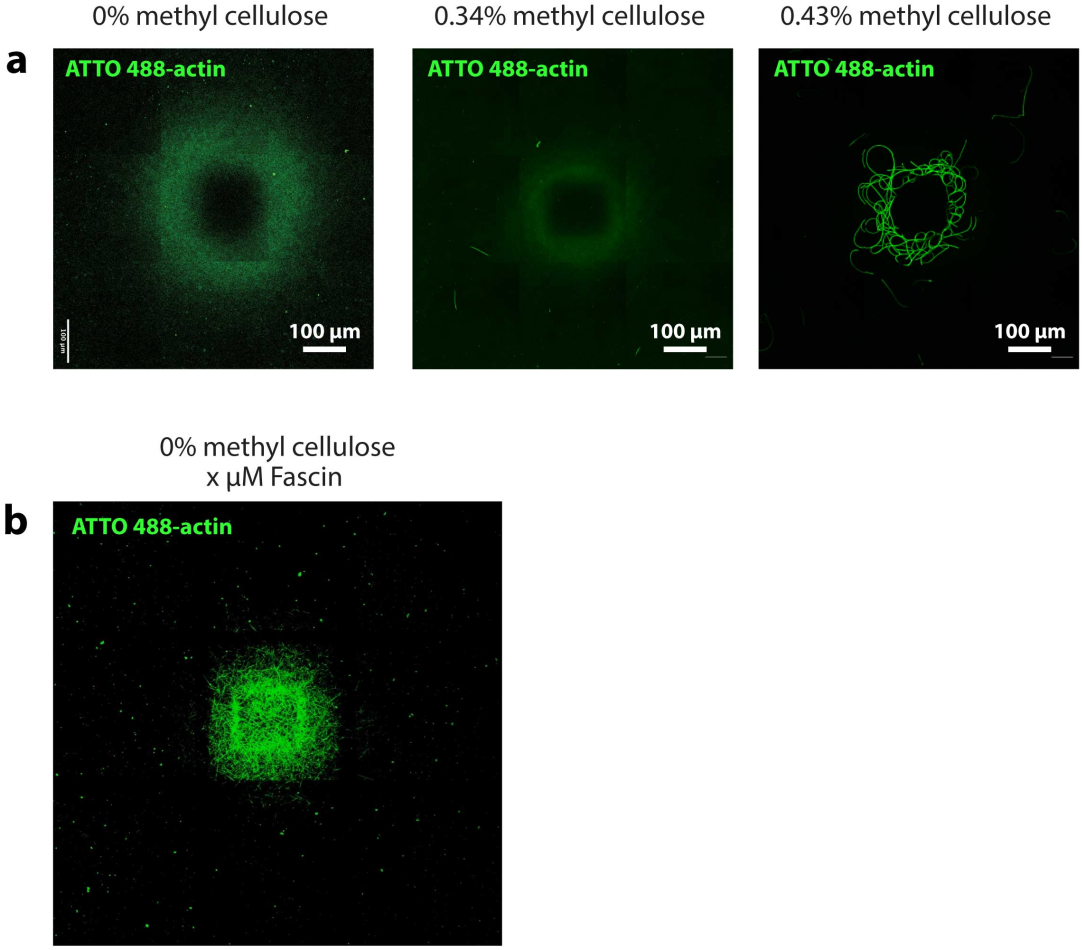


**Figure S1:** Actin printing in dependence on crowder concentration. **a)** Actin printing can be observed in experiments without any crowder (left panel): we still see a donut-shaped pattern even without methyl cellulose in experiments with high laser power, i.e. an increase in F-actin surrounding the exposed region. However crowder (here in the form of methyl cellulose) affects the results. With increasing methyl cellulose, the printing becomes less diffuse, possibly due to a higher viscosity and reduced diffusion. If we increase crowder concentration even further, we cross a bundling transition and actin bundles start to appear. In our experiments bundles usually form close to the laser-exposed region, where actin concentration is highest, although also the opposite can occur (Figure S20). **b)** Bundling in the light-affected area can also be achieved using bundling proteins instead of crowder. Shown is an experiment without any crowder, but with the bundling protein fascin. Similar to conditions with crowder, this limits printing to a smaller area. 1.5 µM actin, 0.5 µM fascin.

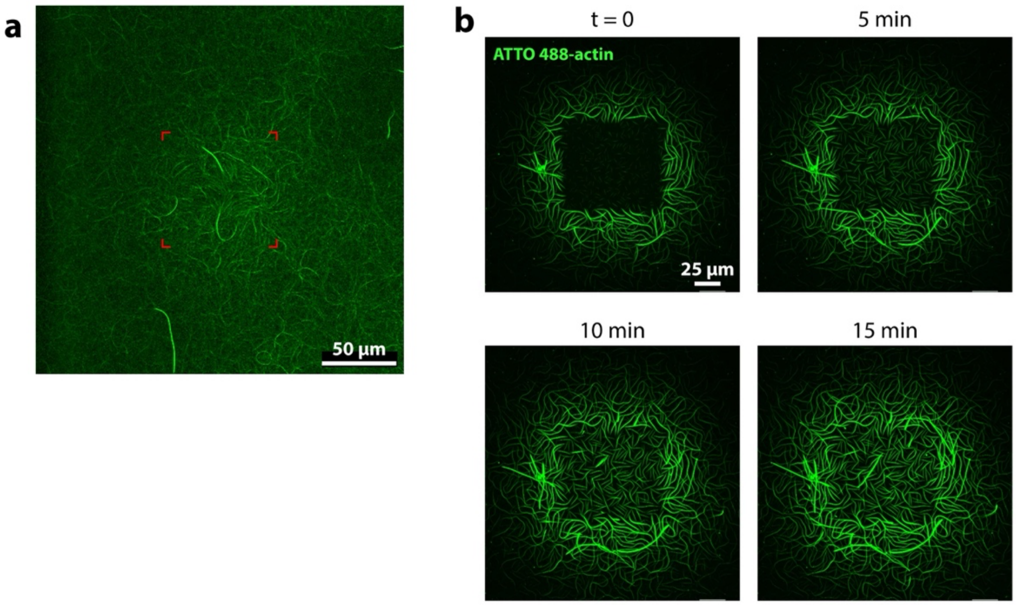


**Figure S2:** Light exposure causes sustained actin assembly beyond illumination period. **a)** Even a very short exposure of 10 seconds leads to an increase in actin. Shown is a sample with 1.5 µM actin and 0.04 µM rose bengal. The samples was left to polymerize for 15 minutes, then the region in the center (red mark) exposed at 514 nm light for 10 seconds and left to polymerize for another 10 minutes. After 10 minutes (shown here) an increase in actin can be seen in the exposed region. **b)** Sample with 1.5 µM ATTO 488-actin. This sample was light exposed for 50 minutes at high light intensity, after which light exposure was stopped and the first image (t = 0 min) was taken. Consecutive images are taken 5 minutes apart without additional light exposure. We observe continued actin assembly in the previously light exposed-region and to some extent in the directly surrounding area (ring).

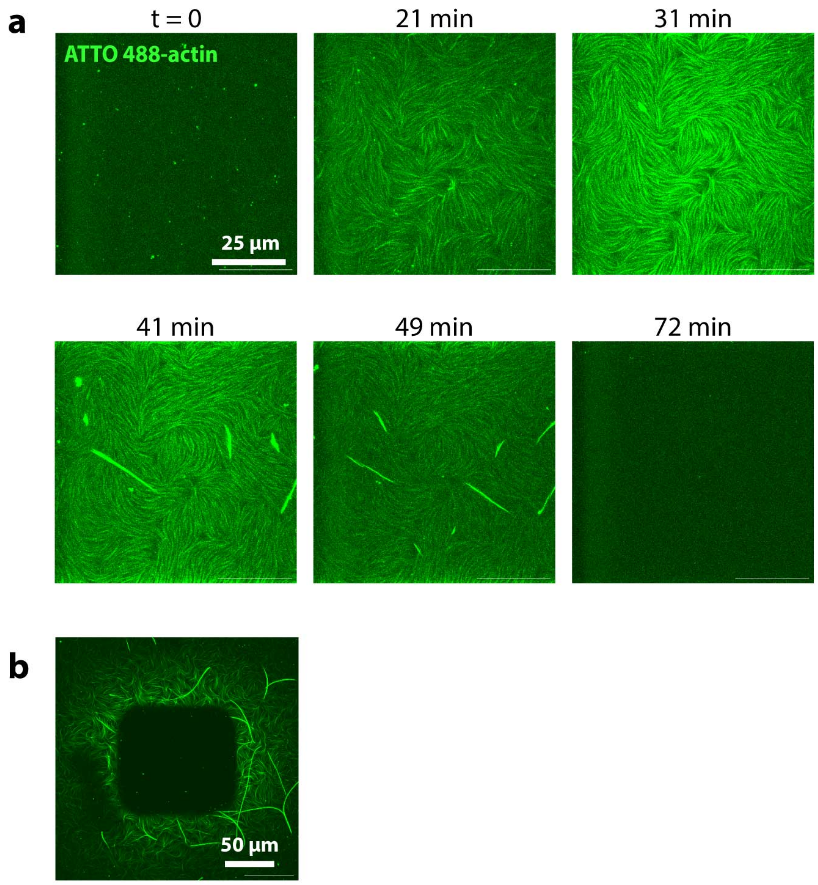


**Figure S3:** Effects of light exposure over time. **a)** Field of view is the immediately light exposed area. Development over the course of 72 min. We see a progression from actin bundles appearing to them disappearing. **b)** Zoomed out view of the area after the light exposure (t = 72 min). 1.5 µM actin.

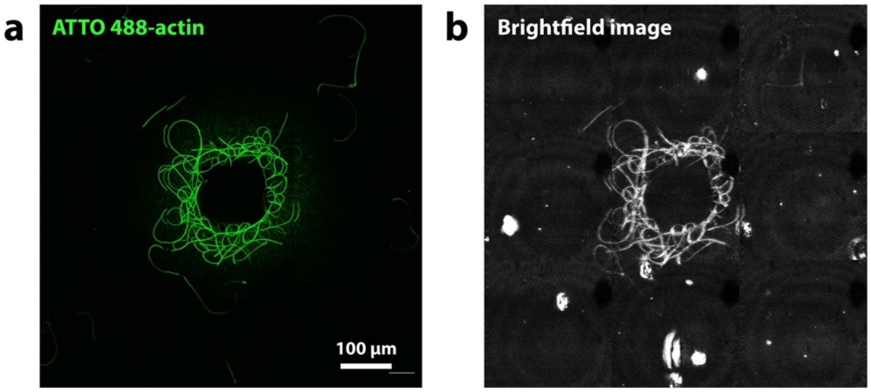


**Figure S4: Comparison to brightfield image. a)** Fluorescence microscopy image and **b)** brightfield image of a ring-shaped print. The brightfield image is heavily processed, which allows us to see thick actin bundles. The similarity of the two images indicates that photo-related artifacts like photobleaching are not a major contributor to the lack of fluorescence in the exposed region.

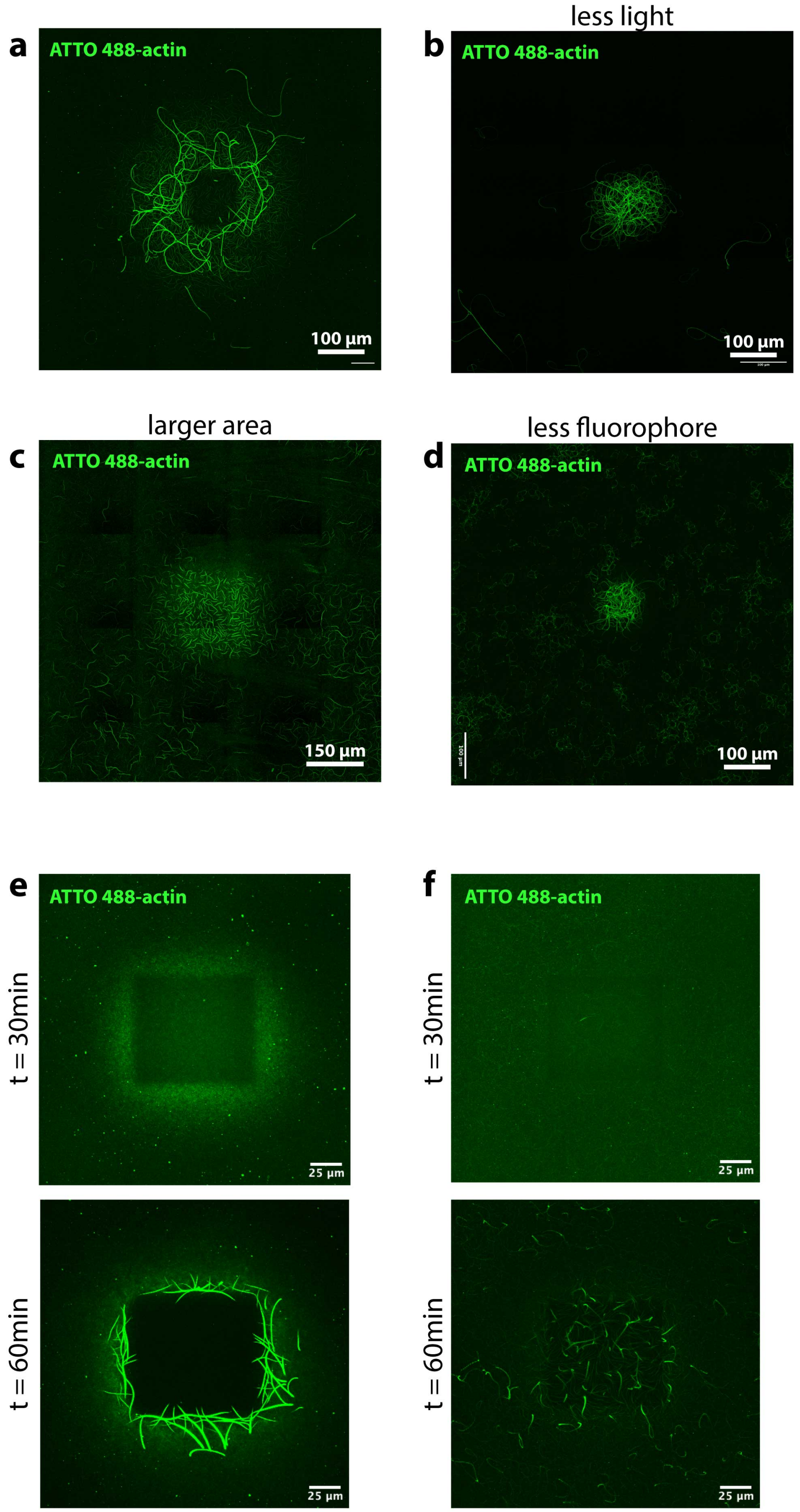


**Figure S5:** With decreasing amount of ROS generation we see a transition from a ring-shaped print pattern with actin disassembly in the center, to a purely additive print. **a)** "Control"

experiment, similar to donut print experiments in Figure 1d. 1.5 μM actin of which 70% are fluorescently labeled (ATTO 488-actin), exposed area is ~100 μm x 100 μm in size. **b)-c)** Experiments in which ROS generation (over space or time) is decreased, result in additive print. Each image shows experiment in which one parameter was adjusted compared to a). **b)** Experiment with less light exposure. Same laser intensity, but only 25% of the time the laser is on; 75% off-time. **c)** A larger area is light exposed, which, using a confocal microscope, results in a lower light density. Exposed area is ~270 μm x 270 μm. **d)** Experiment with less fluorescently labeled actin (23% ATTO 488-actin, 77% unlabeled actin). **e)** and **f)** More detailed comparison between experiment with 70% labeled actin (e)) and 23% labeled actin (f)).

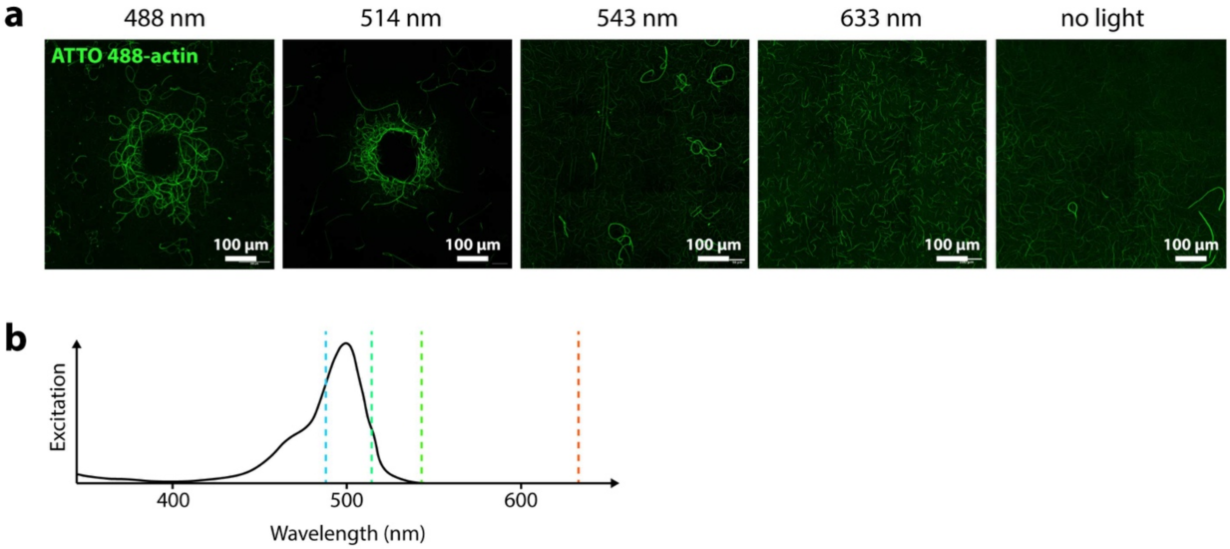


**Figure S6: a)** Actin printing experiments with high laser power and 1.5 µM ATTO 488-actin. Compared are experimental runs in which sample is exposed to lasers of different wavelengths. Printing is only observed for 488 nm and 514 nm, which are laser wavelengths within the excitation spectrum of the fluorophore. **b)** Excitation spectrum of ATTO 488. Adapted from (*AAT Bioquest, Inc. (2024, November 11). Quest Graph™ Spectrum [Atto 488]. AAT Bioquest. https://www.aatbio.com/fluorescence-excitation-emission-spectrum-graph-viewer/atto_488*). Four excitation wavelengths from a) are marked with dotted lines. Normalized absorption for 488 nm is 69%, for 514 nm is 39 % and for 543 nm and 633 nm are both <1%.

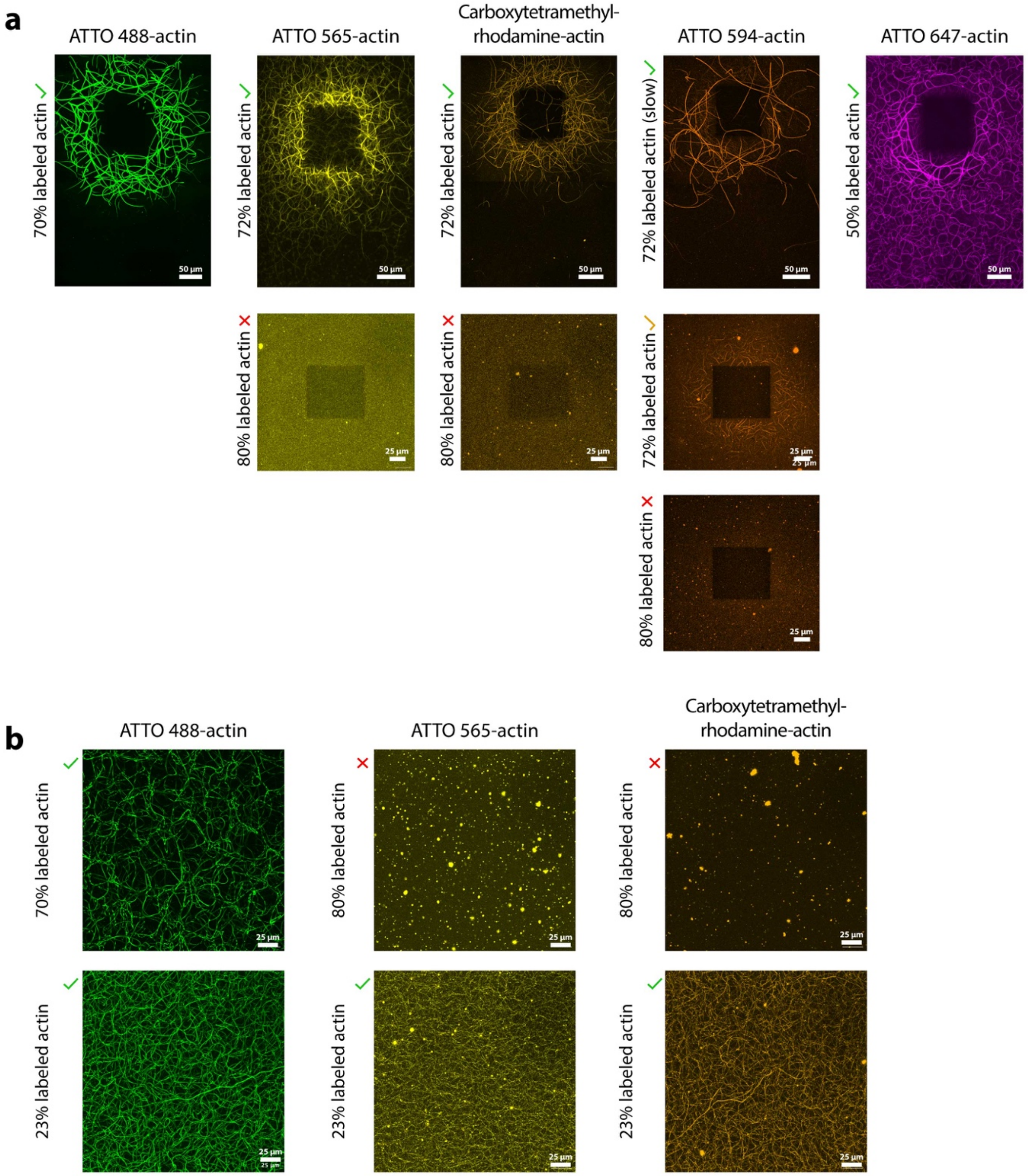


**Figure S7: a)** Different fluorescently labeled actins at various degrees of labeling. At high degrees of labeling, we neither observe actin printing nor other kinds of polymerization (see Figure S7b). Additional information regarding different fluorescent labels: We note that different fluorescent labels behave differently and have to adjust conditions for consistent outcomes. As such we use different total actin concentrations: 1.5 µM for ATTO 488-actin, 1.0 µM for ATTO 565-actin, 1.75 µM for ATTO 594-actin, 1.5 µM for ATTO 647-actin and 1.8 µM for Carboxytetramethylrhodamine-actin. Further, we were only successful with ATTO 594-actin at long exposures (300 min, vs 100 min for all other shown examples here). The highest degree of labeling (most bottom image) for each label shows an experiment with the commercial product in its stock degree of labeling, i.e. not diluted with unlabeled actin. This degree of labeling varies depending on the type of fluorescently labeled actin. **b)** Actin polymerization experiments without light-exposure. Unrelated to ROS-induced printing we note that not all fluorescently labeled actins can be used undiluted at the highest stock ratios of labeled to unlabeled protein.

a

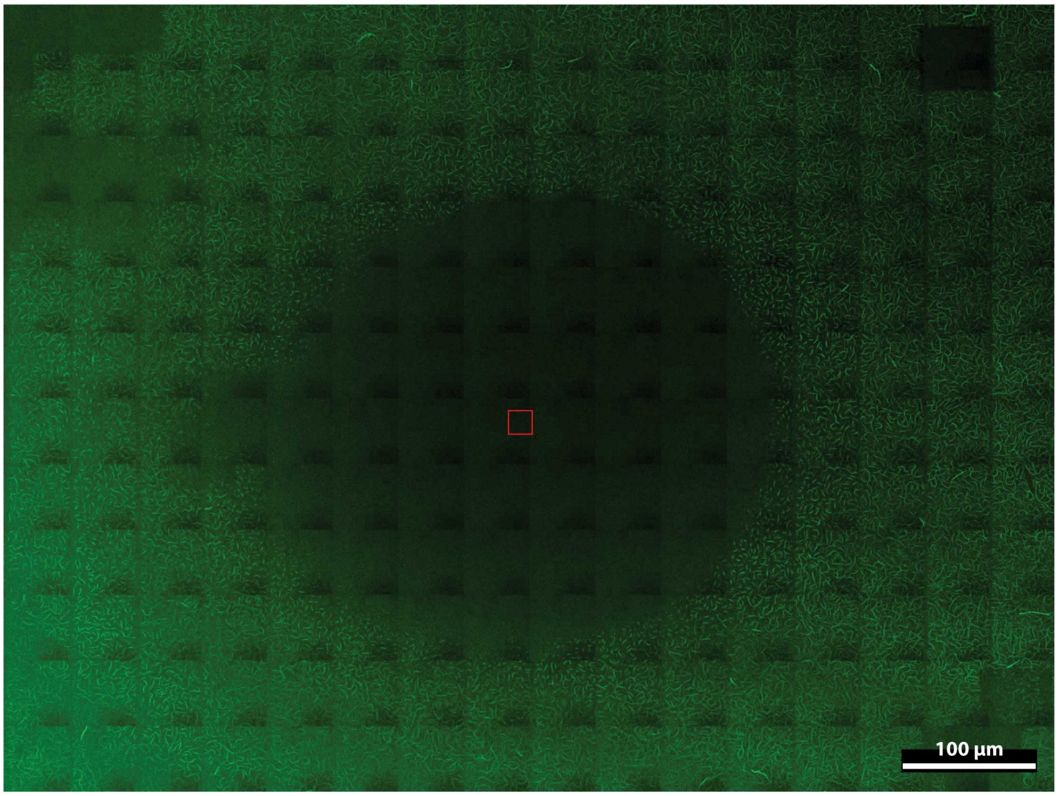


**Figure S8:** Subtractive printing can be diffuse. Experiment with rose bengal, with rose bengal concentrations similar to concentrations of fluorophores in our other experiments. Due to rose bengals propensity to produce ROS very efficiently, ROS is produced in large concentrations in this experiment. This high concentration of ROS is allowed to diffuse further, hence fragmenting actin far away from the light-exposed area.

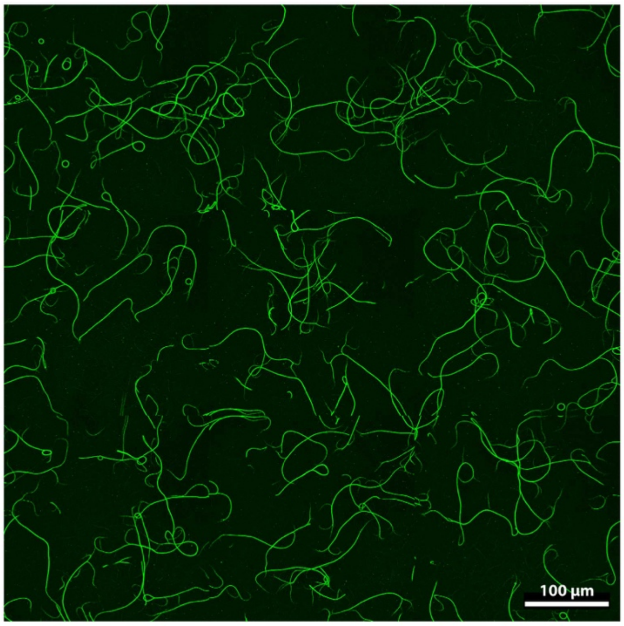


**Figure S9:** Experiment with gold nanoparticles. Gold nanoparticles produce thermal energy when excited within their size-specific excitation wavelength spectrum. We do not see any effect on actin polymerization or disassembly when we excite gold nanoparticles via laser excitation. Experiment with 1.5 µM actin (35% ATTO 488-actin), 0.43% methyl cellulose, 2 mM ATP-γ-s. This sample has with $4.7*10^{12}$ particles/mL of 5 nm gold nanoparticles, which have excitation maximum between 510 nm and 520 nm. Sample was exposed at 514 nm for 150 min.

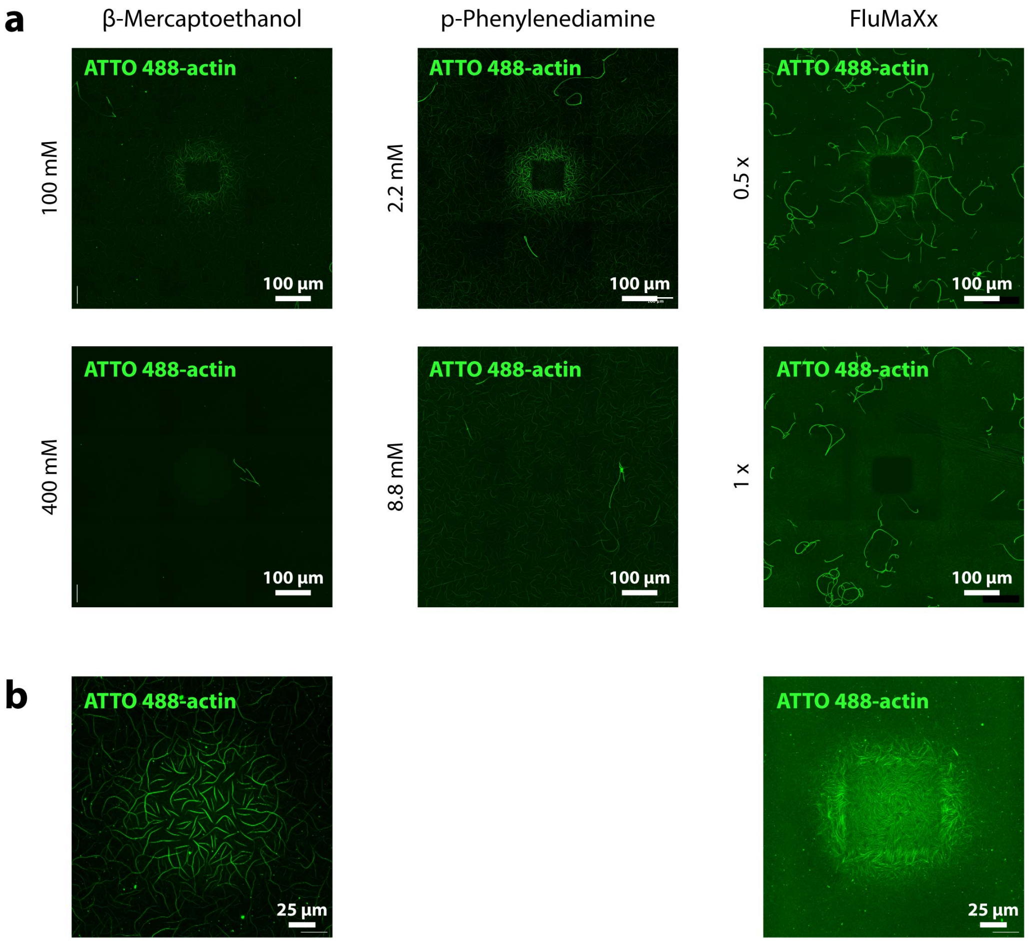


**Figure S10: a)** Experiments with antioxidants and oxygen scavenger. Conditions with 1.5 µM ATTO488-actin, same as experiment shown in Figure 2a, however here with the addition of antioxidants/oxygen scavenger and longer exposure. β-Mercaptoethanol (BME) and p-Phenylenediamine (PPD) are antioxidants. FluMaXx is a commercial oxygen scavenger system consisting of the enzymes glucose oxidase and catalase and glucose as a substrate. Long exposure times (2 – 3 hours) were chosen to test for both additive and subtractive printing. **b)** When exposed for a shorter time, positive printing can be observed, similar to other cases with lower ROS-production rate (see Figure S9). Shown here is an experiment with 100 mM BME (left) and 0.5 x FluMaXx (right) after 45 min and 30 min of exposure respectively. In both cases additive printing is prevalent. Without these compounds (BME and FluMaXx) these exposure times would result in a donut-shaped print (subtractive in the center). The exposed regions are the same size as in a), but note the smaller field of view in both images compared to a).

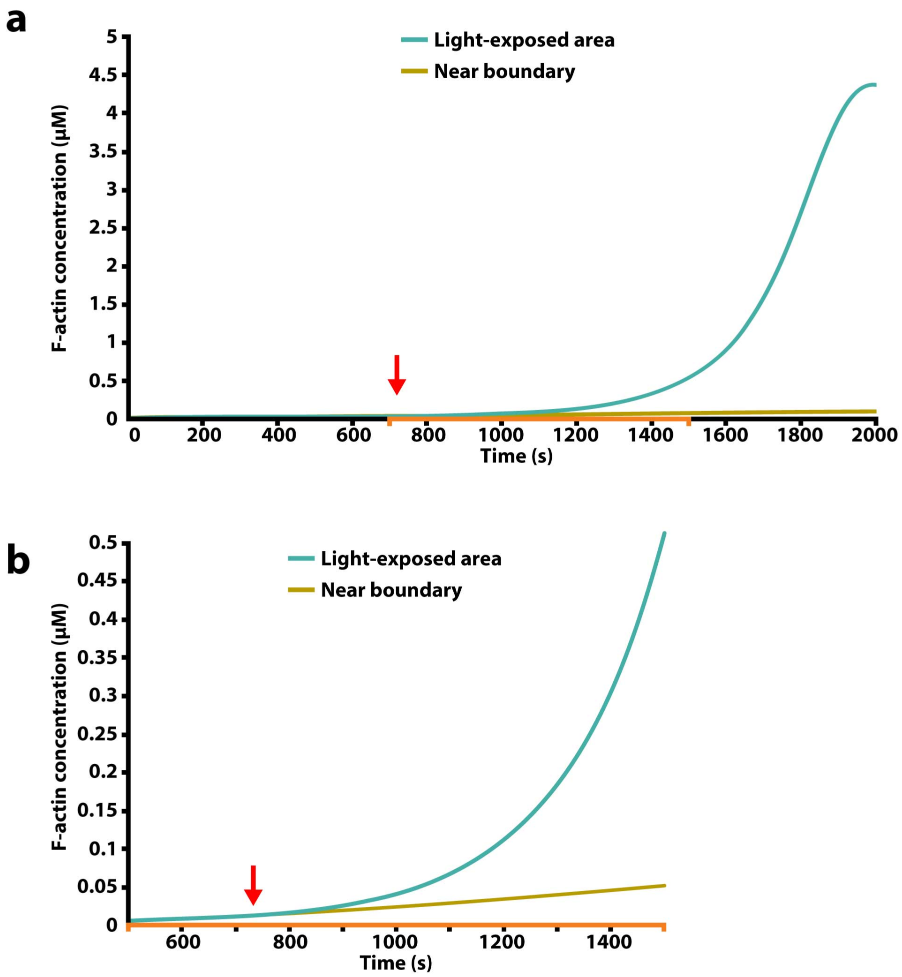


**Figure S11: F-actin concentration over time in simulation of light-increased actin polymerization. a)** Model with low light intensity, exposing an area of 75 x 75 µm in a sample of 225 x 225 µm. Initial G-actin concentration is 0.75 µM. Light exposure starts at t= 720 s (red arrow). Cyan curve shows the F-actin concentration in the very center of the light-exposed region. Yellow curve shows F-actin concentration near the boundary of our simulated sample, which is mostly unaffected by the light-exposure. **b)** Zoom in on time interval between t = 700s and t = 1500s of a). Initial exponential increase in F-actin can be seen, after light-exposure and before G-actin depletion slows down polymerization.

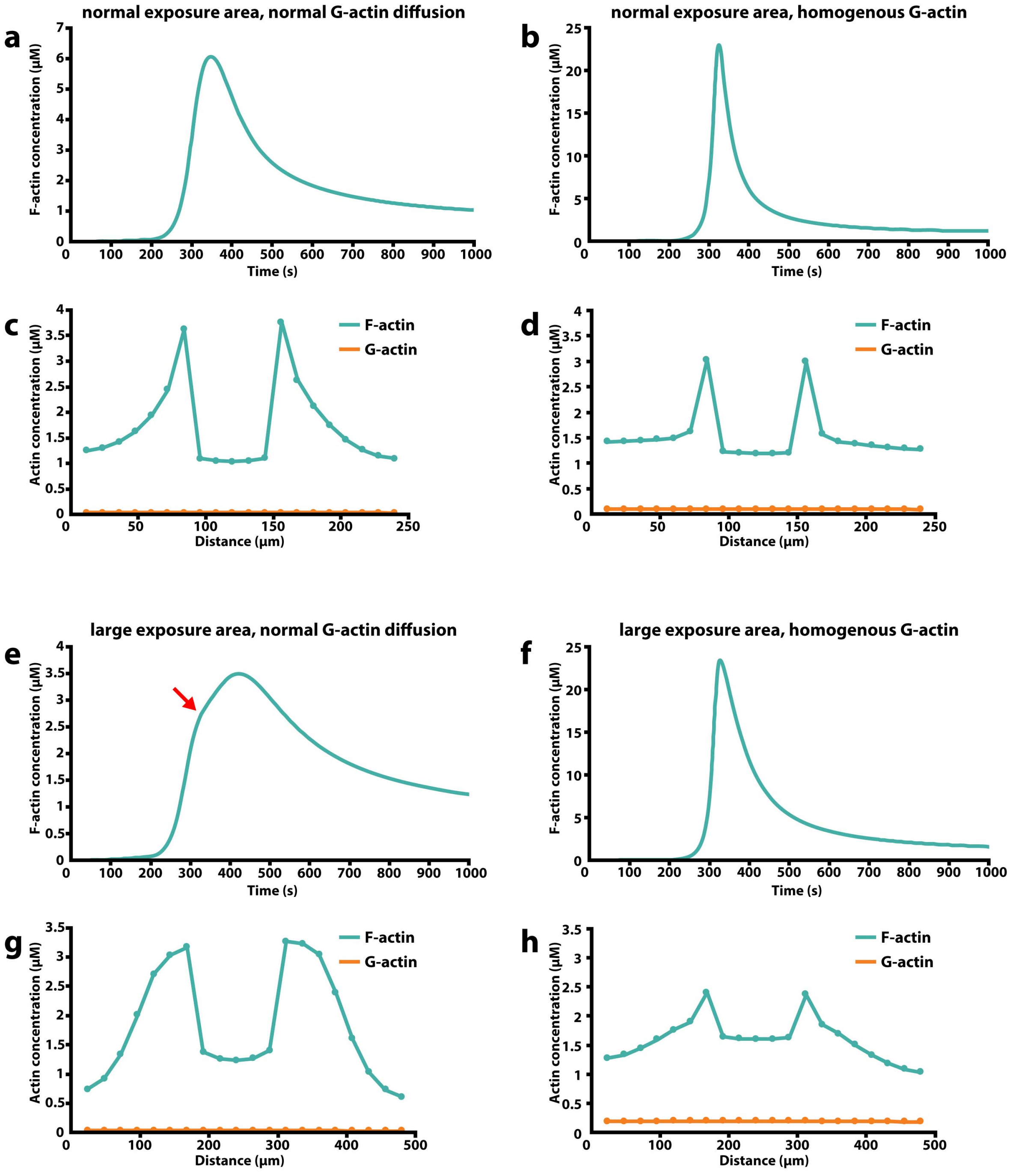


**Figure S12: Effect of G-actin diffusion on printing.** We design a simulated experiment so that G-actin concentration is homogenous across space at all times, essentially simulating a diffusion coefficient that is infinitely high. **a), b), e), f)** show F-actin concentrations over time in the light-exposed region. **c), d), g), h)** show the concentration profile across the sample at t = 1000s for all conditions. All cases are with 1.5 µM actin. The directly light exposed region is visible as a dip in F-actin concentration in each plot. Four simulations are shown. The first column shows two experiments with normal G-actin diffusion. The second column shows two experiments in which we homogenized G-actin concentration across space. The top two simulations show a smaller simulated sample with a smaller light-exposed area. The bottom two simulations show a larger sample with a larger light-exposed area. As expected, the increased G-actin diffusion accelerates actin polymerization, as the pool of G-actin is constantly replenished. Note the vastly different y-axis a scales in figures a), b), e) and f). For smaller exposed areas, the concentration profile at 1000s changes relatively little when comparing the two different G-actin diffusion scenarios [c) and d)]. For larger exposed areas, G-actin diffusion affects the outcome more strongly [g) and h)]. Also note a kink that can be seen in the F-actin concentration curve (red arrow) in g), which shows the complete depletion of available G-actin, which disappears for high G-actin diffusion [f)].

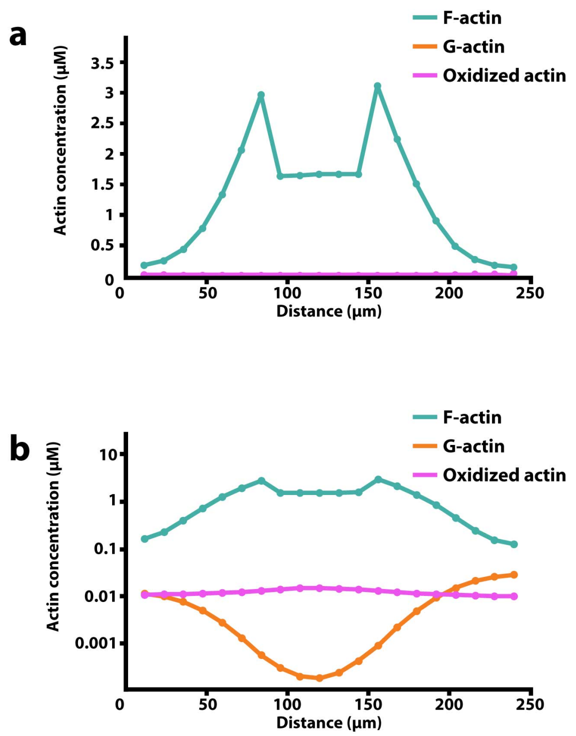


**Figure S13: Actin oxidation.** Simulation with 0.75 µM actin at t = 7200s. **a)** Concentration profile of F-actin, G-actin and oxidized actin across central axis of sample. **b)** Same plot as a), with logarithmic scale. Concentration of oxidized actin monomers is very low compared to total actin concentration.

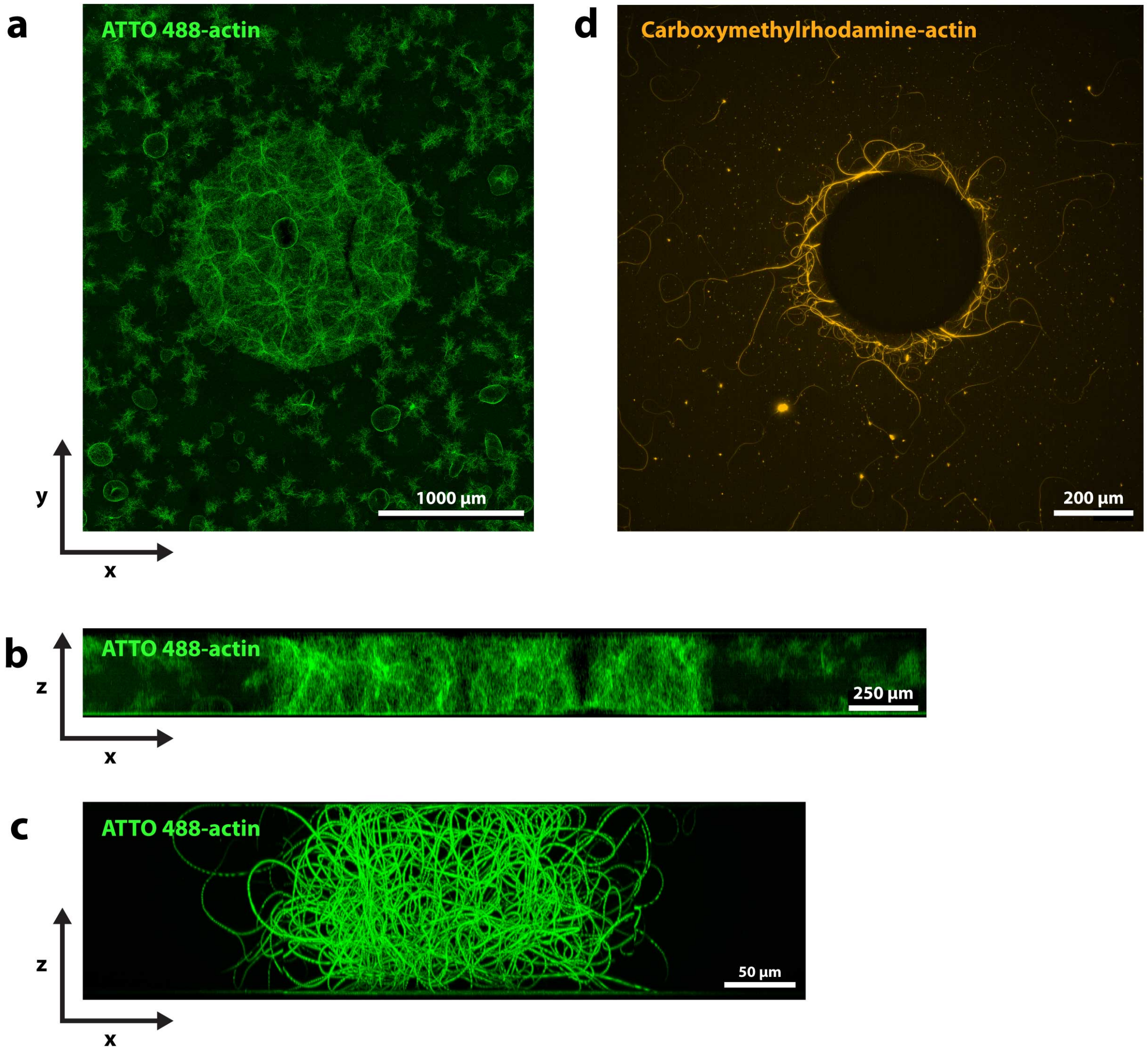


**Figure S14:** Large actin prints. **a)** Actin print similar to Figure 4a, but with a lower actin concentration (1.2 µM), lower fraction of labeled actin (23% ATTO 488-actin) and exposed for a longer duration (10 hours) at low light intensity. **b)** Side view of the sample (maximum projection), shows that the print expands in z, resulting in a large printed volume. We use this sample to run a SDS gel to check for possible oxidation-induced crosslinking of actin monomers (Figure S21) **c)** Shows a large print, printed on a confocal microscope. **d)** Actin print performed on a widefield microscope with carboxymethylrhodamine-actin. Sample was exposed for 5 hours using 475 nm and 555 nm light simultaneously. Sample with 1.7 µM actin (50% carboxymethylrhodamine-actin).

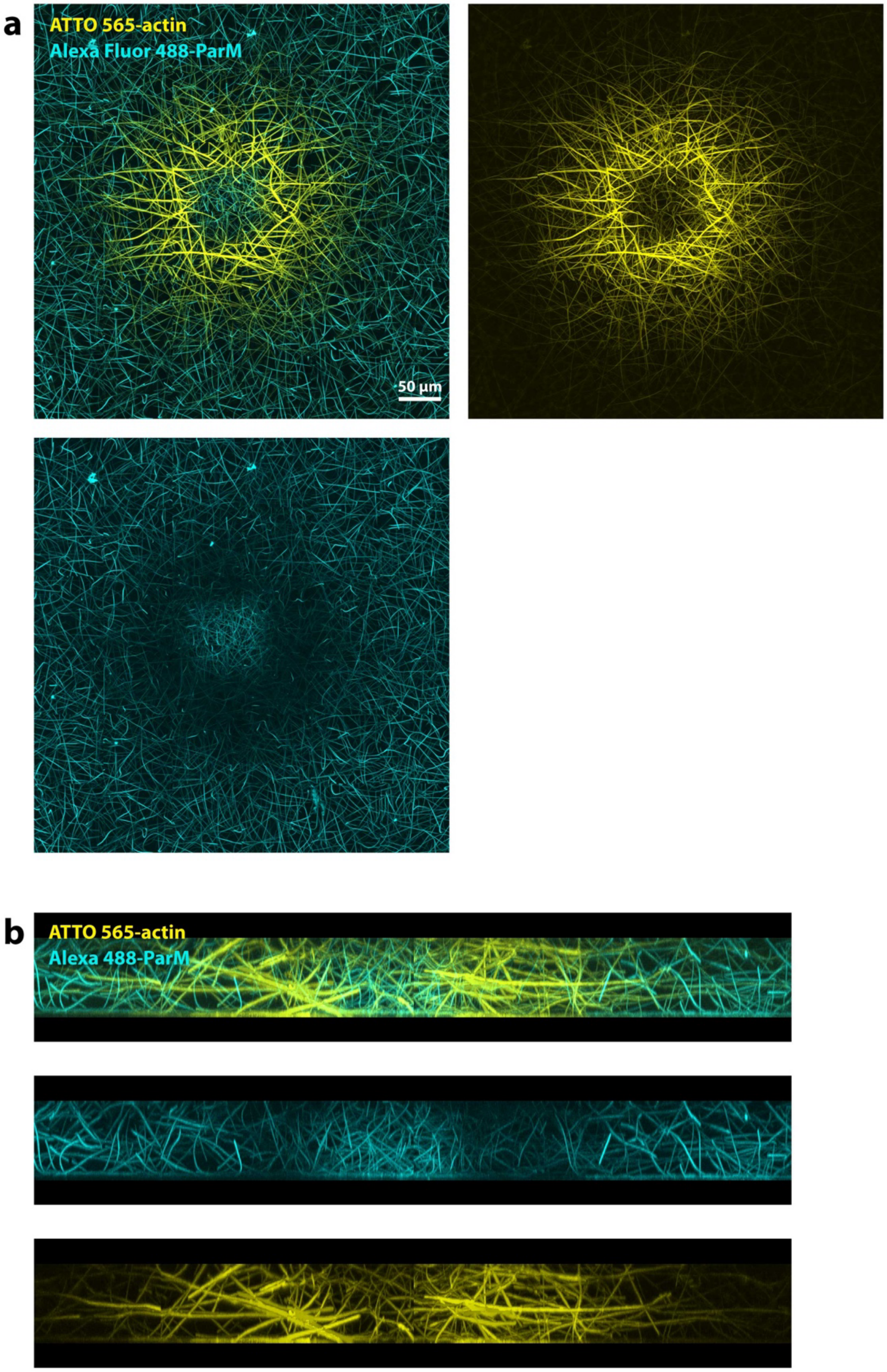


**Figure S15:** ParM-actin multi-color print with 22 µM ParM (35% labeled) and 2 µM Actin (70% labeled). **a)** Similar to the experiment in Figure 5, but with a more pronounced decrease in ParM in region high in actin. **b)** Side view cross-section of the sample. Shown is a maximum projection of the center region (80 µm thick) of the sample. Actin tends to form thicker bundles, which here are mostly horizontally (x-y) oriented, while ParM bundles are thinner and more vertically (z) oriented, which could suggest different material properties in this multi-material print.

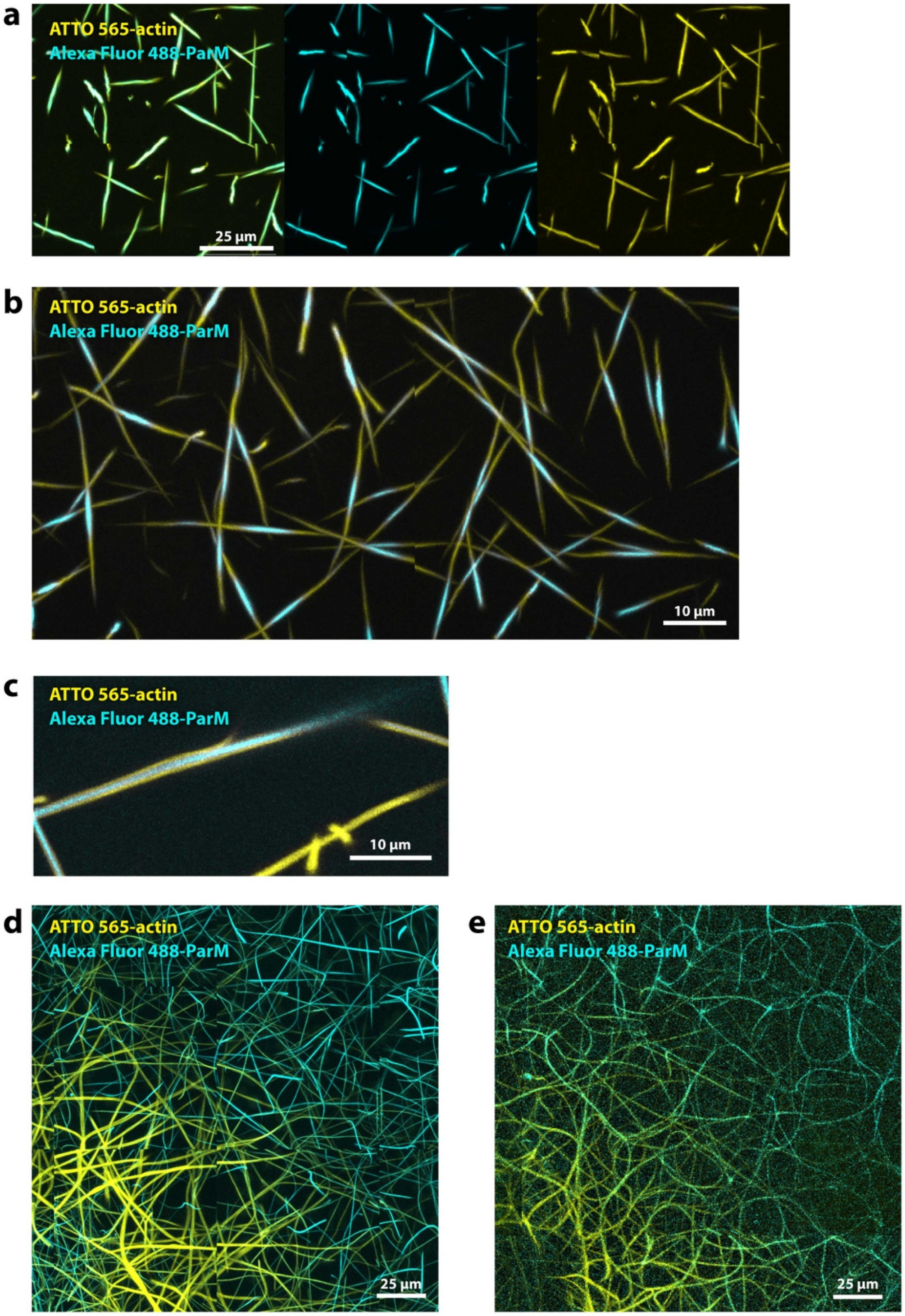


**Figure S16:** Colocalization of actin and ParM. **a)** We often see colocalization of actin and ParM within one bundle. We assume that actin and ParM polymerize independently, i.e. form separate filaments, but form bundles together. **b)**, **c)** Due to the different polymerization kinetics of the two filaments patterns can form within the bundles, both longitudinally (b))

and laterally (c)). **d)**, **e)** ParM and actin "multi-color" prints, zoomed in on the area in which the transition from actin-rich region to ParM-rich region occurs. Usually ParM and actin do not co-localize within bundles (d)), however in some cases the transition happens within bundles (e)).

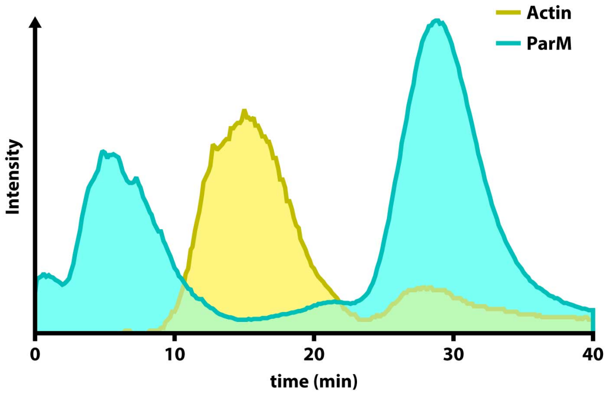


**Figure S17: Actin and ParM fluorescence intensity over time.** Plotted is the fluorescence intensity over time of the entire field of view of Movie 3. Interestingly we see three phases, where first, ParM assembles in the light-exposed region, second ParM disassembles while actin assembles, and third, actin disassembles while new ParM bundles form.

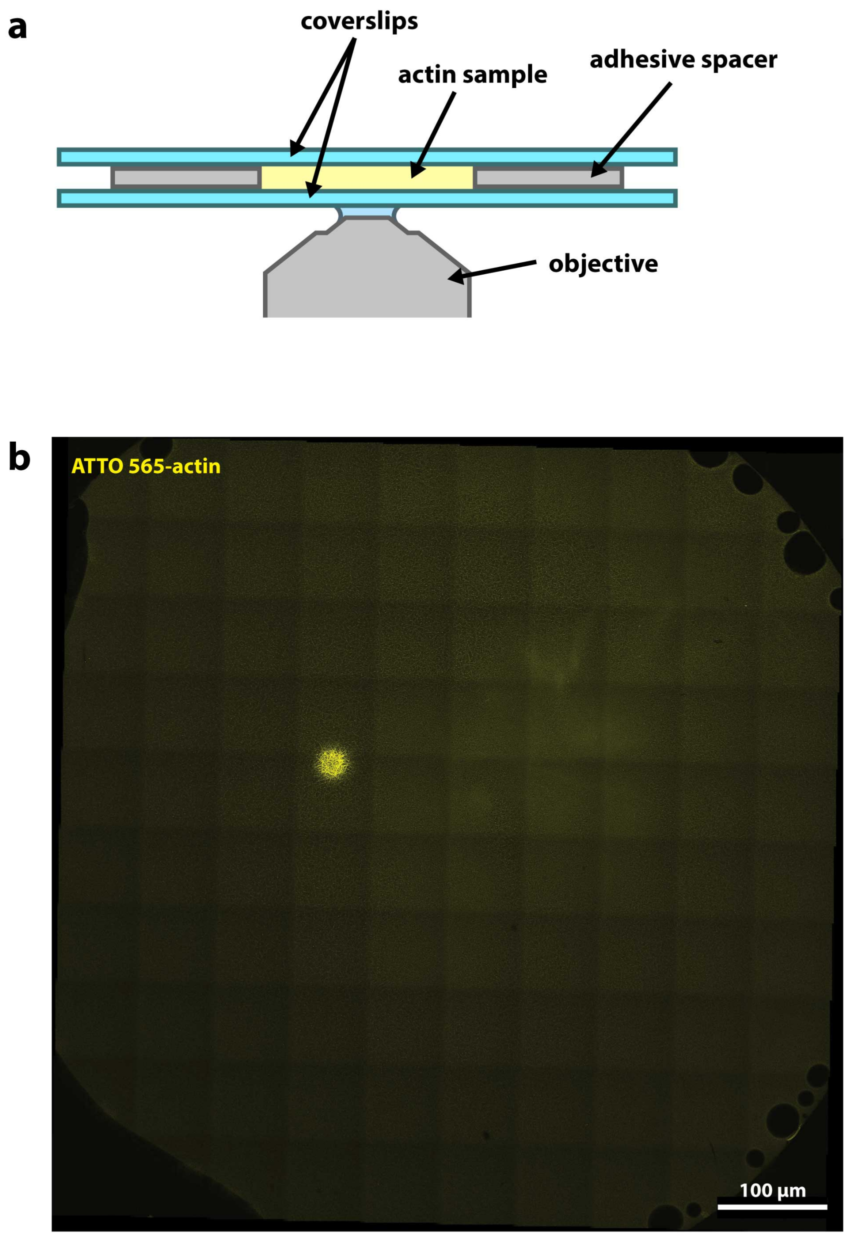


**Figure S18: Experimental setup. a)** Illustration of side-view of the setup. We use two 1 mm thick coverslips with a 200 µm thick spacer with a circular cutout in between the coverslips. **b)** Microscopy image that shows an entire sample. Tile scan stitched together from 100 individual images. Towards the center of the image a light exposed area can be seen with an increase in actin. 2 µM actin.

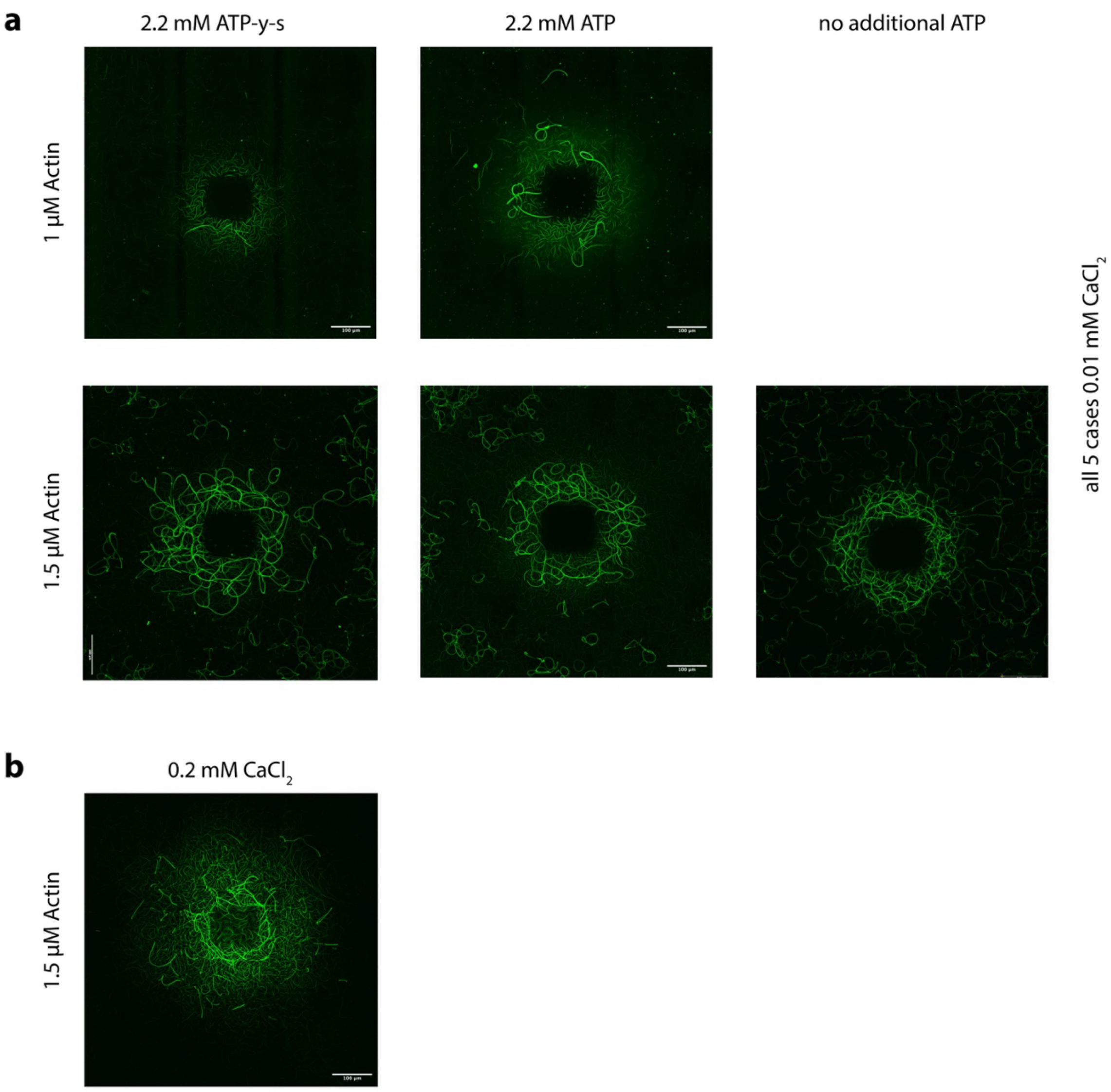


**Figure S19:** Comparison with more common polymerization conditions. We acknowledge that two parameters in our polymerization buffer deviate from conditions most commonly used in actin polymerization experiments. In control experiments we do not see a significant difference when compared to commonly used conditions. **a)** Unless otherwise noted, in all of our experiments we use ATP-γ-s, a slowly hydrolyzing variant of ATP. A comparison between ATP and ATP-γ-s shows no significant difference between the two types of ATP. It should be noted that we use regular ATP in our G-actin buffer in which our actin is stored. As such, even in our experiments with ATP-γ-s, likely most actin is bound to ATP during polymerization and not ATP-γ-s. In fact, we see similar outcomes if we do not add any additional ATP or ATP-γ-s in our F-buffer in experiments (right image). **b)** Typical polymerization buffers contain higher concentrations of $CaCl_2$ than we use. However, we see no significant difference in this control experiment with increased $CaCl_2$ concentrations.

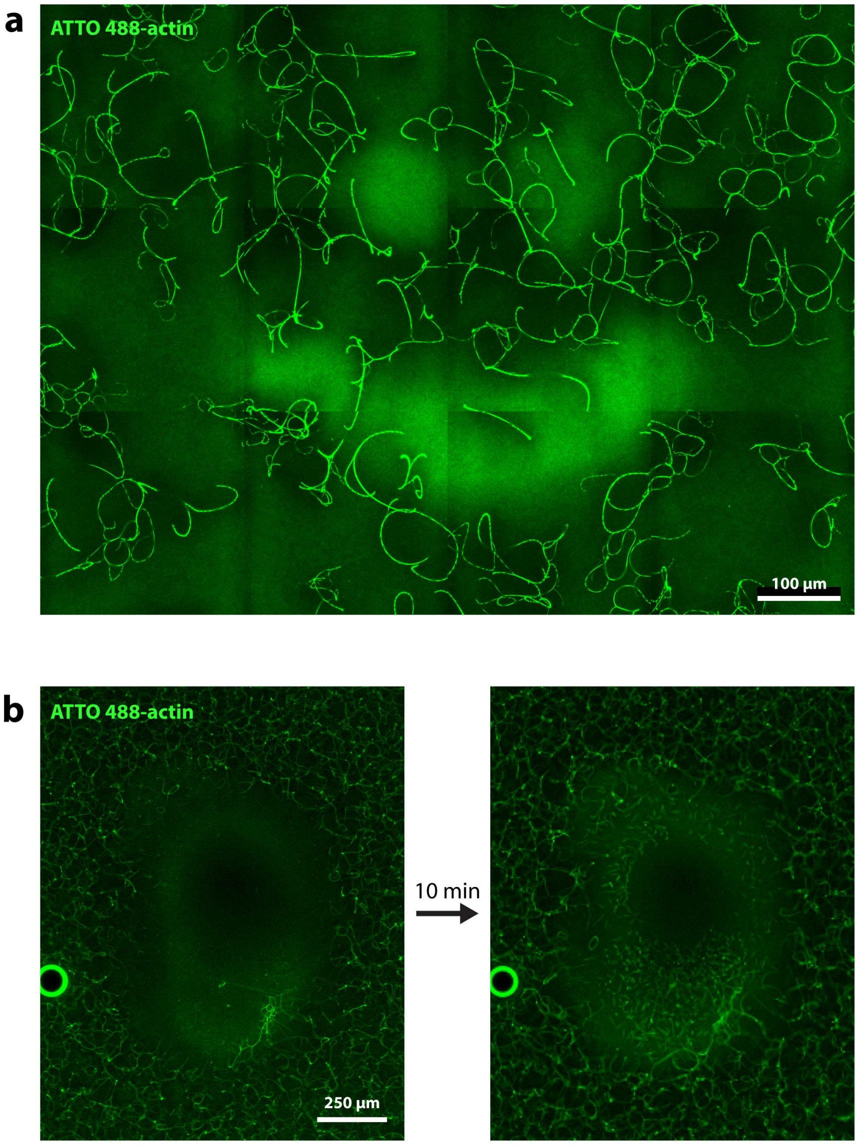


**Figure S20:** Light exposure can prohibit bundling due to filament severing. **a)** Light exposure in smiley shape (see Figure 4d), but with higher initial actin concentration (2 µM) and stronger laser exposure. This lead to a print in which we see an increase in F-actin in the light exposed area, however, without the formation of bundles. At the same time, we do see bundles in the periphery, due to the increased initial actin concentration. We think this indicates that the bundles are shorter in the exposed area compared to the periphery. Bundling both depends on the concentration of filaments and their length; it seems here we found a condition in which bundles in the exposed area are so short, that despite an increased filament concentration, bundling does not set in. **b)** Similar observation as in a). Experiment performed on a widefield microscope, also at higher actin concentration (2 µM). We exposed part of the sample for 15 minutes (oval shape in the center) and see disassembly of the filaments, but also an increase in actin signal. After 10 minutes without light exposure, short, but thick bundles start appearing.

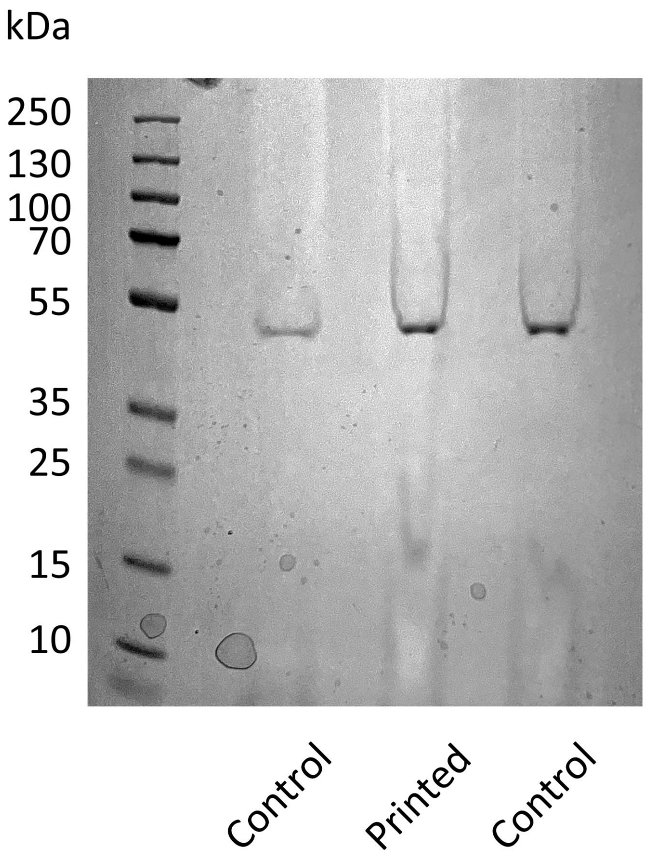


**Figure S21: SDS page gel of printed sample.** We test for oxidation-induced crosslinking of actin monomers, which we do not detect. The sample which is shown in Figure S14a,b was used for this gel, as the printed volume was particularly large.
